# Tree diversity reduces pathogen damage in temperate forests: a systematic review and meta-analysis

**DOI:** 10.1101/2024.09.16.613230

**Authors:** Elsa Field, Andrew Hector, Nadia Barsoum, Julia Koricheva

**Author notes:** Corresponding author: Andrew Hector.

## Abstract

Diversifying planted forests to reduce the risks associated with large scale disturbances, such as pathogens, is a major aim of sustainable forest management. Previous meta-analyses have shown that insect pest damage is lower in mixed forest stands compared to monocultures, but the same has not been shown for pathogens. Here, we provide the first systematic review and meta-analysis specifically testing the effects of tree species diversity on pathogen damage. Relevant studies were retrieved using bibliographic databases and internet searches, as well as previously unpublished data sets contributed by stakeholders. We found that more diverse forest stands overall had significantly lower pathogen damage, and that this result was most pronounced in temperate forests for which the most studies were available. Although in some cases tree diversity had a strong effect, this was not universal and was influenced neither by pathogen specialism, nor by the presence of alternative hosts in stands. Instead, we found that tree neighbour identity rather than species richness emerged as a crucial variable impacting pathogen damage in mixed stands. Neighbour identity effects reported in studies were far-ranging, including impacts on the microclimate of stands. Future work should focus on mechanistic explanations that could underpin neighbour identity effects in mixed forests. We suggest the use of the disease triangle as a tool for considering the multiple factors that can influence pathogen damage in mixed forest stands.

## 1 Introduction

Plant pests and pathogens are a major threat to forest health worldwide (Fisher et al., 2012). Even small, chronic loss of leaf tissue due to herbivory has been shown on a global scale to lower forest productivity by 2-15% (Zvereva et al., 2012). As well as reducing ecosystem services provided by trees, plant pests and pathogens cause significant economic costs to the timber industry every year (Haight et al., 2011; Boyd et al., 2013). There is a clear need for durable strategies to reduce infection transmission and improve overall forest productivity and resilience to disease (Ennos, 2015). Planting mixed species stands is often promoted as a way to reduce pest and pathogen damage and thus increase forest productivity (Forrester & Bauhus, 2016; Jactel et al., 2017).

The negative relationship between plant diversity and plant diseases has been widely demonstrated in agricultural systems and experimental grassland communities (Mitchell, Tilman, & Groth, 2002; Mundt, 2005). Disease theory states that as the density of a given host reduces in mixture, there is lower potential for pathogen populations to multiply, disperse and cause disease – the dilution effect (Keesing et al., 2010; Mundt, 2005). A meta-analysis by Civitello et al. (2015) provided evidence of the dilution effect occurring across diverse study systems ranging from animals to plants. This study included just one paper on forests. Likewise, Liu et al. (2020), in a meta-analysis of plant diversity effects on disease, found a reduction in pathogen damage overall in mixtures, which was nonsignificant in forests. However, this study included only six papers considering pathogens on trees, five from Europe and one from Mexico. Roberts et al. (2020) provided an evidence synthesis of forest management effects on pathogens, including tree species diversity, which suggested that negative relationships between diversity and pathogens were most common. However, whether tree diversity positively or negatively impacts pathogen damage has yet to be the focus of a comprehensive, global meta-analysis or systematic review.

It has been suggested that effects of plant diversity on pathogen infection could be smaller in forests compared to other plant communities because the large size of trees may lead to increased frequency of autoinfection (multiple infections on the same host plant), reducing the effectiveness of mixtures (Mundt, 2002). Furthermore, given the comparatively long generation time of most forest plantations, the ability of a mixed stand to avoid and resist disease progression over the entire length of the rotation has to be questioned.

In contrast to forest pathogens, there is strong evidence that tree diversity has a negative impact on specialist insect herbivores. This phenomenon is known as associational resistance (Castagneyrol et al., 2014; Jactel & Brockerhoff, 2007; Vehviläinen et al., 2007). However, when the tree species in a mixture are close in phylogenetic relationship to one another, tree diversity effects on insect pest damage are reduced (Castagneyrol et al., 2014). In fact, when neighbouring hosts are more susceptible to insect pests than a focal host, associational susceptibility can occur, where spillover occurs onto the focal host from more susceptible hosts, reversing the protective effect of mixed stands on pest damage (Barbosa et al., 2009). These studies suggest that both pest specialism and tree neighbour identity play important roles in determining tree diversity effects on insects.

We lack an understanding of whether associational resistance in mixed forests also occurs for forest pathogens. As trees are often under threat from multiple biotic factors simultaneously, this makes determining forest diversity effects on pathogens even more important (Poeydebat et al., 2021; Jactel et al., 2017; Manion & Lachance, 1992). A literature review of biodiversity effects on pathogens in natural communities found that neutral/positive relationships between host diversity and pathogen damage were just as common as negative relationships for non-specialist pathogens (van der Plas, 2019). The same study found that for specialist pathogens, negative relationships outweighed positive/neutral ones (van der Plas, 2019). This suggests that host specialism may play an important role for pathogens as it does for insects, although the study did not quantify this using meta-analysis. Previous work in mixed forests also found tree species composition effects as well as diversity effects on pathogens (Hantsch et al., 2014; Hantsch et al., 2013). The evidence on the different factors which impact upon host diversity-pathogen damage relationships in forests has yet to be assimilated.

In this study, we focused on forests and undertook a global systematic review and meta-analysis of the English language forestry literature relating to mixed forest stands and pathogens. We included both experimental and observational studies from across the globe and searched both bibliographic databases and the grey literature. We tested the following specific hypotheses in the meta-analysis:

1) Overall, mixed species forest stands would have lower pathogen damage than monocultures, consistent with evidence for a dilution effect.
2) There would be a stronger negative effect of tree species mixtures on specialist pathogens than generalist pathogens.
3) The presence of alternative hosts in mixed stands would lead to either reduced damage on focal hosts or an increase in damage, depending on whether alternative hosts were more or less susceptible than the focal host. Alternative hosts being less susceptible may increase host dilution, leading to associational resistance. Alternatively, the presence of more competent hosts might lead to an associational susceptibility effect.

In addition to these more general hypotheses, we also summarised results from studies containing qualitative information and observations on the topic and focused on extracting information on mechanisms that could underpin diversity-disease effects.

## 2 Methods

We followed the established protocol for systematic reviews from the Collaboration for Environmental Evidence (Pullin & Stewart, 2006). Our systematic review protocol was independently peer-reviewed blind by a panel of experts recruited through the CEE network (by handling editor Neal Haddaway). Comments on the methodology were synthesised and the resulting protocol was submitted to Research Gate to be available open access before data collection began. We summarise the methodology used here, and further details are available in the protocol (Field, Petrokofsky and Koricheva, 2018).

### 2.1 Literature searches and inclusion criteria

We used the PECO framework (components of the primary question, comprising population, exposure, comparator and outcome) as the basis for study searches and study inclusion criteria. To be judged relevant to the review, a study needed to include all four components. We identified the *population* as tree species, including both long and short rotation crops and fruit trees. The *exposure* (treatment) was mixed tree species stands, the *comparator* (control) was single species stands, and the *outcomes* were metrics of pathogen damage (incidence, severity and abundance measures) on the focal tree species. We included both observational / longitudinal studies from natural (unplanted) forests, and experimental studies from tree diversity experiments and forest plantations. We only included studies where the English full text of the paper was available. No restrictions were given on the publication date of studies.

We searched three major bibliographic databases: Scopus, Web of Science (relevant databases only, listed in Appendix I) and CAB Abstracts. We also searched Google Scholar, as Haddaway et al. (2015) showed only poor overlap between results from Web of Science and Google Scholar when similar search strings were used. We first conducted a scoping search using Scopus, refining search terms based upon whether they could return articles in the test library of studies known to be relevant (Appendices I-II). The final search terms used included a range of keywords relevant to each component of the primary question (Table S1). Details of searches conducted and numbers of hits in each database are given in Table S2.

A range of stakeholders known to be working in the field were also contacted and asked to contribute relevant studies, as well as the personal libraries of contributors (Appendix III). We also searched the citations of articles found to be relevant at the level of full text, to find further studies.

We also received unpublished data from TreeDivNet, the global network of forest diversity experiments (Verheyen et al., 2016). Data was contributed along with standardised metadata sheets, submitted by each contributor (Appendix IV), and these underwent the same process of screening as for all other studies.

The total number of studies brought forward to the screening stage was 11,470 (Fig. 1).

**Figure 1:**
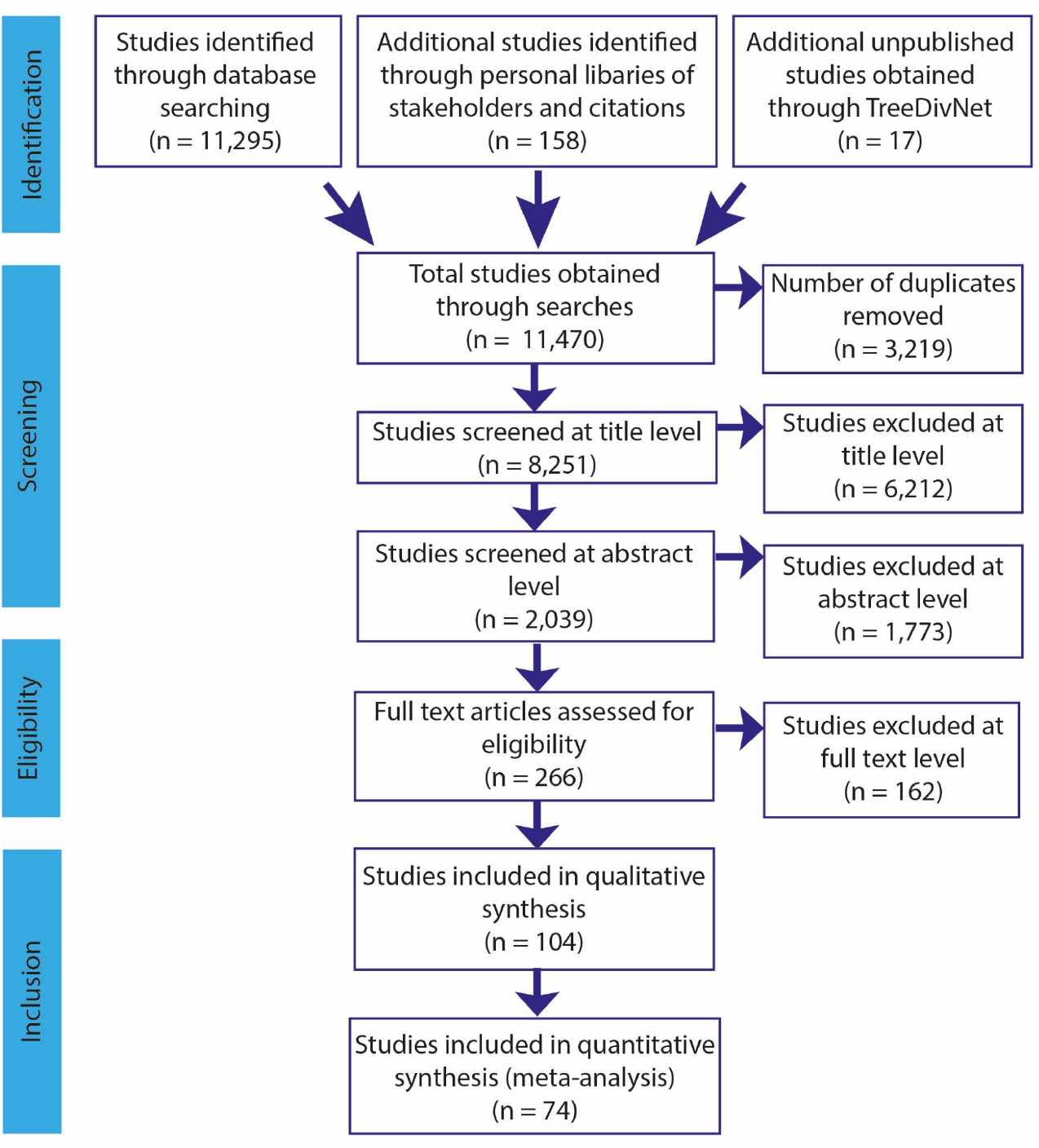
PRISMA record workflow of studies included in the systematic review. Divided into sections: Identification of studies, Screening of studies, Eligibility of studies, Inclusion of studies. The number of studies at each stage is indicated in brackets by *n*.

### 2.2 Article screening

Studies were assessed for their relevance for inclusion first based on title, then title and abstract, then full text (Fig. 1). Screening at both title level and abstract level was completed using the online tool Abstrackr: (http://abstrackr.cebm.brown.edu/). The majority of article screening was undertaken by the same reviewer (EF) but we performed kappa tests at both title and abstract screening stages to check for consistency. All reviewers received the same list of inclusion criteria for assessing studies. A kappa value > 0.6 indicated that reviewers were consistent in their assessments of study relevance. If k < 0.6, then reviewers discussed the possible sources of inconsistency and repeated until k > 0.6. A randomised subset of 200 articles were used for kappa tests at both title and abstract level.

### 2.3 Data Extraction

We used standardised spreadsheets for extracting all data from publications (Field, Petrokosky & Koricheva, 2018). First, we assessed whether a publication contained sufficient quantitative data for inclusion in the meta-analysis. If so, then publication metadata and quantitative data were extracted, along with any additional qualitative data. Where data were available only in graphs, we used the online tool Web Plot Digitizer (https://automeris.io/WebPlotDigitizer/) to extract data. The same reviewer (EF) carried out data extraction for all studies. If insufficient quantitative data was available for the meta-analysis, we extracted metadata and qualitative data only.

We used Standardised Mean Difference (Hedges’ *d*) as the effect size to test for differences in pathogen damage between monospecific and mixed species stands (Hedges, 1981). We chose to use Hedges’ *d* as the measure of effect size, as:

1. Many studies included comparisons between monocultures and just one level of multispecies mixtures (for example, two-species mixtures), rather than continuous relationships between tree species richness and pathogen damage.
2. It is possible to convert to *d* to/from other statistics that may be available from studies (such as *t-*statistics and correlation coefficients) using standard equations (Koricheva et al., 2013)
3. Hedges’ *d* has commonly been used in similar meta-analyses assessing plant diversity effects on plant attackers, which enhances the comparability of our study with others in the field (e.g. Jactel & Brockerhoff, 2007; Koricheva & Hayes, 2018).

To calculate Hedges’ *d* we extracted means, sample size and standard deviation of treatment and control groups from each publication. A positive effect size indicated that pathogen damage was higher in mixed species stands compared to monocultures, while a negative effect size indicated that mixed species stands had a protective effect.

In some cases, multiple effect sizes could be calculated from the same article, i.e. the same article was split into multiple studies. This was the case for instance if damage by multiple pathogen species was measured in the same article, or if a paper contained data from multiple study sites that had not been pooled.

Wherever possible, we calculated effect sizes directly from data available in the study. For studies where a continuous relationship between more than one level of tree diversity had been compared to monoculture, we used standard equations to convert between correlation coefficients and Hedges’ *d* (Koricheva et al., 2013). If insufficient data was available to calculate effect size directly, we firstly contacted the authors for additional data. If additional data were unavailable, we obtained Hedges’ *d* and its variance by interconversion between effect sizes from the data available in the study using standard equations (Koricheva et al., 2013). In the case of unpublished studies from TreeDivNet, we calculated effect sizes directly from the raw data provided by authors rather than from summary statistics.

We extracted the following metadata from studies when available in text, tables or graphs.

● Study type (primary research or review - reviews were included in the qualitative evidence synthesis only)
● Study year
● Study geographical location: continent, country, region
● Latitude and longitude of study site
● Biome (boreal, temperate or tropical)
● Size of plots / stands (m^2^)
● Average age of trees at time of sampling
● Focal tree species on which pathogen recorded
● Other tree species present in mixed stands
● Type of focal species (broadleaved or conifer)
● Type of mixture (broadleaved, conifer or mixed i.e. containing both broadleaved and conifer species)
● Pathogen species recorded on focal trees
● Pathogen type (fungal, bacterial, viral or oomycete)
● Pathogen specialism (generalist or specialist, where specialist pathogens were defined as being able to colonise only hosts in the same genera as the focal host)
● Pathogen life history (biotrophic, necrotrophic or both/unknown if multiple pathogens recorded)
● Type of pathogen damage recorded. This fell into three categories: incidence, abundance or severity of pathogen damage depending on how the study was undertaken. Incidence studies described observations of disease presence or absence on trees or in plots, or proportion presence in plots. Abundance studies measured the abundance of infections per plant or plot e.g. the number of lesions per leaf or tree. Severity studies were measures of severity of disease on individual trees e.g. percentage dieback of the crown or percentage coverage by fungal mycelium on leaves.
● Experimental versus observational studies
● Whether summary estimates for pathogen damage were recorded at the level of single trees, or at plot level

### 2.4 Critical Appraisal of Studies

Studies eligible for inclusion at full text were subjected to a critical appraisal of study validity. Studies were assigned into two groups: low or high risk of bias. As there were no geographical or temporal limitations on the review, the critical appraisal focused on internal validity (risk of bias) of studies, rather than external validity (relevance to the review). The same reviewer (EF) carried out critical appraisal of all studies.

Studies were assigned a score based upon the following 2 categories:

1. Presence of treatment replication: use of blocking to account for spatial variability OR presence of multiple study sites: Both (2) / Either (1) / Neither (0)
2. Randomisation: randomisation of tree selection for assessing pathogen damage: Y (1) / N (0)

Studies were then split into two categories – high risk of bias (score of 0-1) or low risk of bias (score of 2-3).

### 2.5 Statistical Analyses

#### Meta-analysis

The total number of studies available for the meta-analysis was *k =* 74 studies, from 44 articles. We used a random effects meta-analysis model as significant heterogeneity between studies was expected. A restricted maximum likelihood approach was used to estimate the parameters of the meta-analysis model. All meta-analyses were conducted using the *metafor* package (Viechtbauer, 2010) in R version 3.5.3 (R Core Team, 2020). We calculated the total heterogeneity in effect sizes (*Q*_t_) as well as the I^2^ statistic, which estimates the proportion of heterogeneity that is present due to between-study variance. Due to the potential non-independence of studies coming from the same paper, we included study ID as a random effect in the meta-analysis (random effect with 44 levels, as per Nakagawa et al., 2017). We used the rma.mv() function in metafor, which enables fitting of multi-level meta-analytic models.

We first tested for the overall significance of the effect of mixed species stands on pathogen damage. Next, we used meta-regression to investigate the impact of categorical moderators on between-study variance. We analysed pathogen specialism and presence of alternative hosts as additional covariates in models, due to our *a priori* hypotheses of interest. We also ran exploratory analyses to test for effects of several other categorical variables that could be extracted from most papers: biome, pathogen life history, type of diversity metric, single tree versus plot level studies, and experimental versus observational studies.

Location information varied per study and we were only able to extract the exact latitude for *k =* 43 studies. Latitude has shown to be a driver of species diversity effects on both pests (Kambach et al., 2016) and pathogens (Liu et al., 2020) previously. Biome was the only location variable available for all studies, and likely included an overall effect of latitude in combination with other environmental factors. For instance, most studies of conifers occurred in the boreal biome (25/30 studies of conifers), while broadleaves were mainly studied in the temperate biome (36/43 studies of broadleaves).

Rather than include moderators together in the same meta-regression model, we chose to fit separate models testing the individual impact of each moderator on effect size.

This was for two reasons:

(1) Not all studies had data available on all moderators, so fitting a model with all moderators would have excluded several studies (reducing the sample size from 74 to 49 studies for a model with all 4 moderators) potentially leading to a biased subsample of studies and reduced statistical power.
(2) Using contingency tables, we found that moderators tended to be confounded. For instance, biotrophic pathogens tended to be specialist pathogens in our data set (Fig. S1). Generalist pathogens tended to be associated with studies where alternative hosts were present in mixed stands. This makes sense, as the likelihood of an alternative host being present in mixed stands will be higher for generalist pathogens (Fig. S1). Specialism was also confounded with biome; more specialist pathogens were present in temperate versus boreal and tropical biomes (Fig. S1).

With several confounded categorical variables, we focused instead on which variable could explain the most variation in effect size on its own, potentially giving us a clue towards underlying mechanisms. For the model including Pathogen Specialism as a moderator, we excluded studies that tested the effect of both necrotrophic and biotrophic pathogens in the same study (*k* = 6 studies), to provide a clear contrast between necrotrophs and biotrophs only. For the model including Biome as a covariate, we removed studies that took place across multiple biomes (across both Boreal and Temperate biomes, *k =* 3 studies).

#### Sensitivity analysis

Following the main analysis, we conducted sensitivity analyses to assess the sensitivity of the overall effect size to aspects of study design that we had previously assumed did not influence the outcome. To test the effect of study bias on effect size, we ran a meta-regression of overall effect size, including risk of bias score (low or high) as a moderator. We also tested whether conversion between effect sizes had an influence on outcome using meta-regression (i.e., whether or not the metric of effect size (Hedges’ *d*) had been calculated directly or by interconversion from other effect sizes).

#### Publication bias

We tested for potential publication bias using funnel plots, which display effect size plotted against standard error, and inspected them for visual asymmetry. Potential publication bias is indicated by asymmetry in the plot, with studies reporting small or nonsignificant effects missing from the opposite side of the plot to the true effect size. However, funnel plots have limitations, as asymmetry can be caused by several other factors (Lau et al., 2006). We tested for the significance of asymmetry of the funnel plot using Egger’s regression (function *regtest()* in *metafor*). We also calculated fail safe numbers using Rosenberg’s weighted method to assess the robustness of results to potential publication bias (Rosenberg, 2005). The failsafe number indicates the number of additional studies having an effect size of 0 that would be required to add in order to make the overall effect size statistically non-significant (Rosenberg, 2005).

## 3 Results

### 3.1 Overall patterns

Our meta-analysis of the effect of mixed species stands on pathogen damage was based on 44 papers yielding 74 comparisons of the effect of tree species mixtures compared to monocultures. We also extracted qualitative data on 30 studies from a further 28 papers.

The majority of quantitative studies (40 out of 74 studies) took place in the temperate biome, 26 were from boreal regions and 5 from tropical regions. This pattern was similar for studies containing only qualitative data, of which 16 out of 30 studies were from temperate regions, 8 from boreal and 4 from tropical regions, with 2 studies spread across both temperate and boreal countries. Fig. 2 shows the patterns of study distribution by geographical country (several studies contained data from more than one country). Quantitative data was concentrated in Europe (with 77 observations in different countries), with studies mostly concentrated in Germany and Finland (19 and 17 studies, respectively). Across other continents, North America had 11 studies in total, South America two studies, Asia had one study (from China), and Africa one study. In studies containing only qualitative data (with 38 observations in different countries), there were 20 studies from Europe, 10 in the USA and the remaining 8 studies between Asia, Africa and South America.

**Figure 2:**
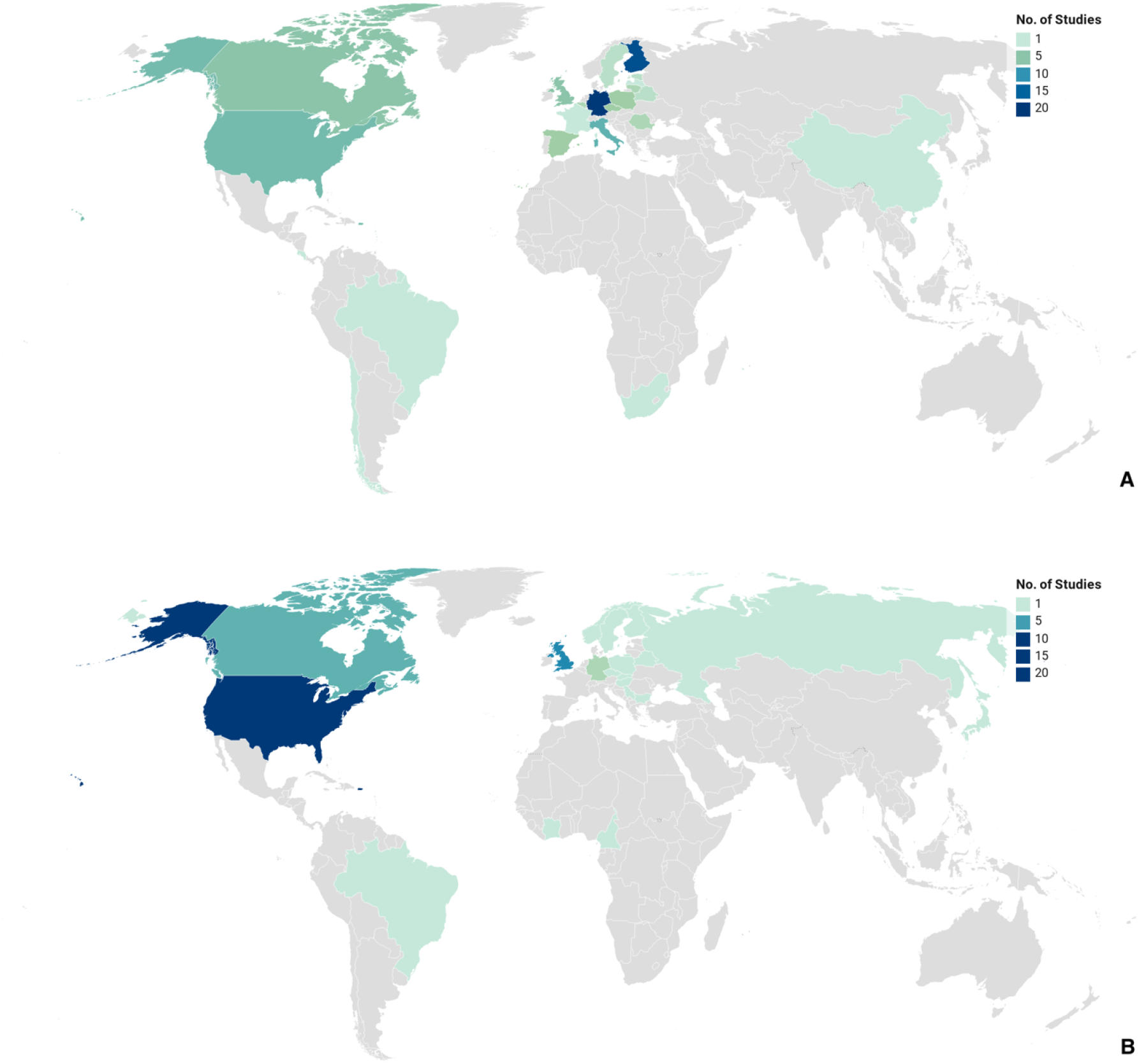
Geographical distribution of countries for which quantitative or qualitative data was extracted. Grey regions are countries with zero studies. Some studies contained data from more than one country. **A)** Quantitative studies only, *n = 93* observations in different countries, total no. of studies (*k)* = 74. **B)** Qualitative studies only, *n =* 38 observations in different countries, total no. of studies (*k*) = 30.

Fungal pathogens were by far the most commonly studied group, making up 68 out of 74 comparisons for the meta-analysis (92%) and 26 out of 30 qualitative studies (87%). The remaining 4 qualitative studies were concerning oomycetes (all *Phytophthora*). One study in the meta-analysis concerned bacteria, three concerned oomycetes (all *Phytophthora ramorum*) and two assessed general pathogen damage symptoms (Fig. 3).

**Figure 3:**
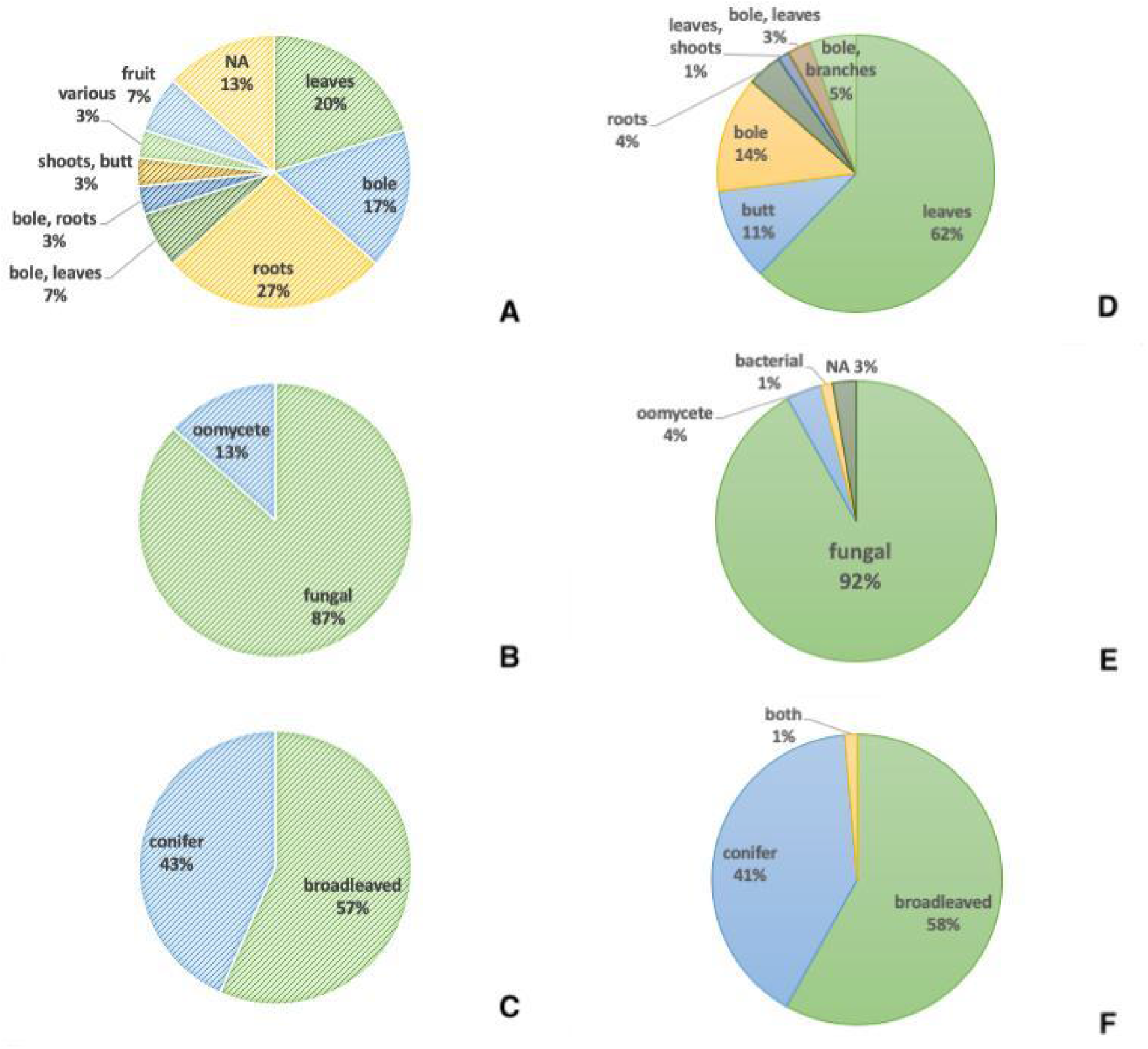
Pie charts showing the distribution of studies by damage organ, pathogen type and host type. **A-C)** Qualitative studies. **D-F)** Quantitative Studies. Explanation of terms used: Butt = the base of the tree stem. Bole = the main portion of the tree stem.

Hosts were divided between broadleaved and coniferous trees. Among quantitative studies, 43 considered broadleaved hosts, 30 considered coniferous hosts, and one study was of both broadleaved and coniferous hosts combined (Ampoorter et al., 2019).

Among qualitative studies, 17 considered broadleaved hosts and 13 considered coniferous hosts (Fig. 3).

Among quantitative studies, pathogens were most commonly studied on leaves, whereas for qualitative studies, work was more evenly distributed between the different organs of the host trees (Fig. 3).

For quantitative studies, the most commonly studied pathogens were rusts and powdery mildews (29 out of 74 studies), in particular the powdery mildew pathogens *Erysiphe* on *Quercus,* which were the focal pathogens of 9 studies. Root rot pathogens were another major group studied; 19 out of 74 studies were of root rot pathogens, all of which were reported on coniferous species.

### 3.2 Meta-analysis

Overall, we found that mixed species stands had lower pathogen damage compared to monocultures (*d =* −0.27, 95% CI −0.49 to −0.046, *k* = 74 studies, Fig. 4). According to Cohen’s original benchmarks for the interpretation of *d* (Cohen, 1970) this corresponds to a small effect size. Between-study heterogeneity was high (Q_t_ = 835.67, *p* < 0.001, *df* = 73), and 98% of heterogeneity was due to between study variance. This therefore justified the inclusion of additional explanatory moderators using meta-regression.

**Figure 4.**
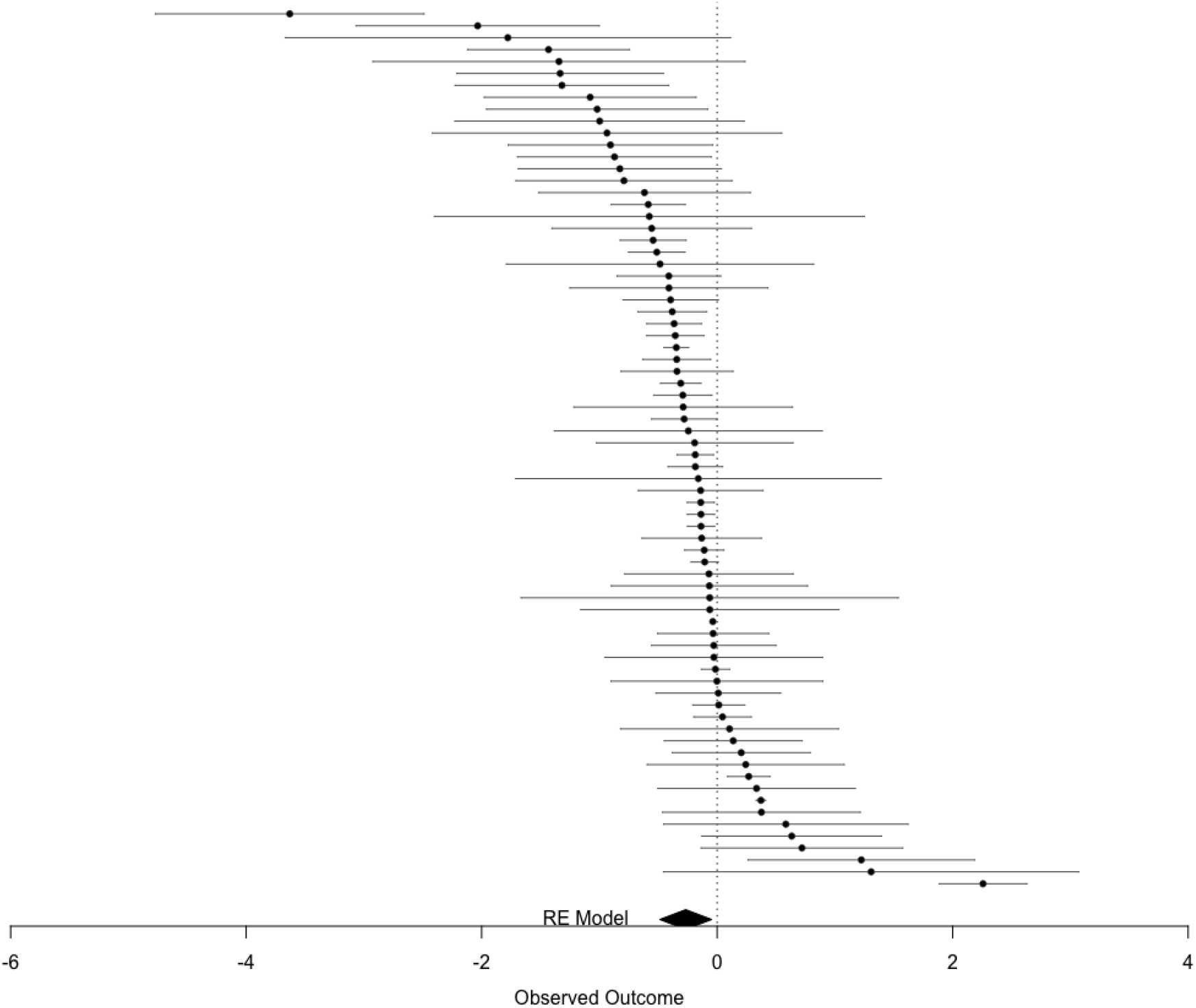
Caterpillar plot showing effect sizes and associated 95% confidence intervals, in order of effect size for all 74 studies included in the meta-analysis. The x axis displays effect size with the vertical dashed line indicating an effect size of 0. Effect sizes are significantly different from 0 if their confidence intervals do not include zero. The bottom polygon shows the overall result from the random effects model (*d =* −0.27, 95% CI −0.49 to −0.046, *k* = 74 studies).

The effect of mixed species stands differed depending on whether the study took place in a boreal, temperate or tropical region (Q_m_ for Biome = 13.17, *p* = 0.0014, *df* = 2, *k* = 71 studies, Fig. 5). Significant reduction in pathogen damage in mixed stands compared to monocultures was observed only in temperate forests, whereas no significant effects of forest diversity on pathogen damage were detected in boreal forests. Results of studies in tropical forests varied but tended to indicate reduction of pathogen damage in mixed stands similarly to temperate forests (Fig. 5).

**Figure 5:**
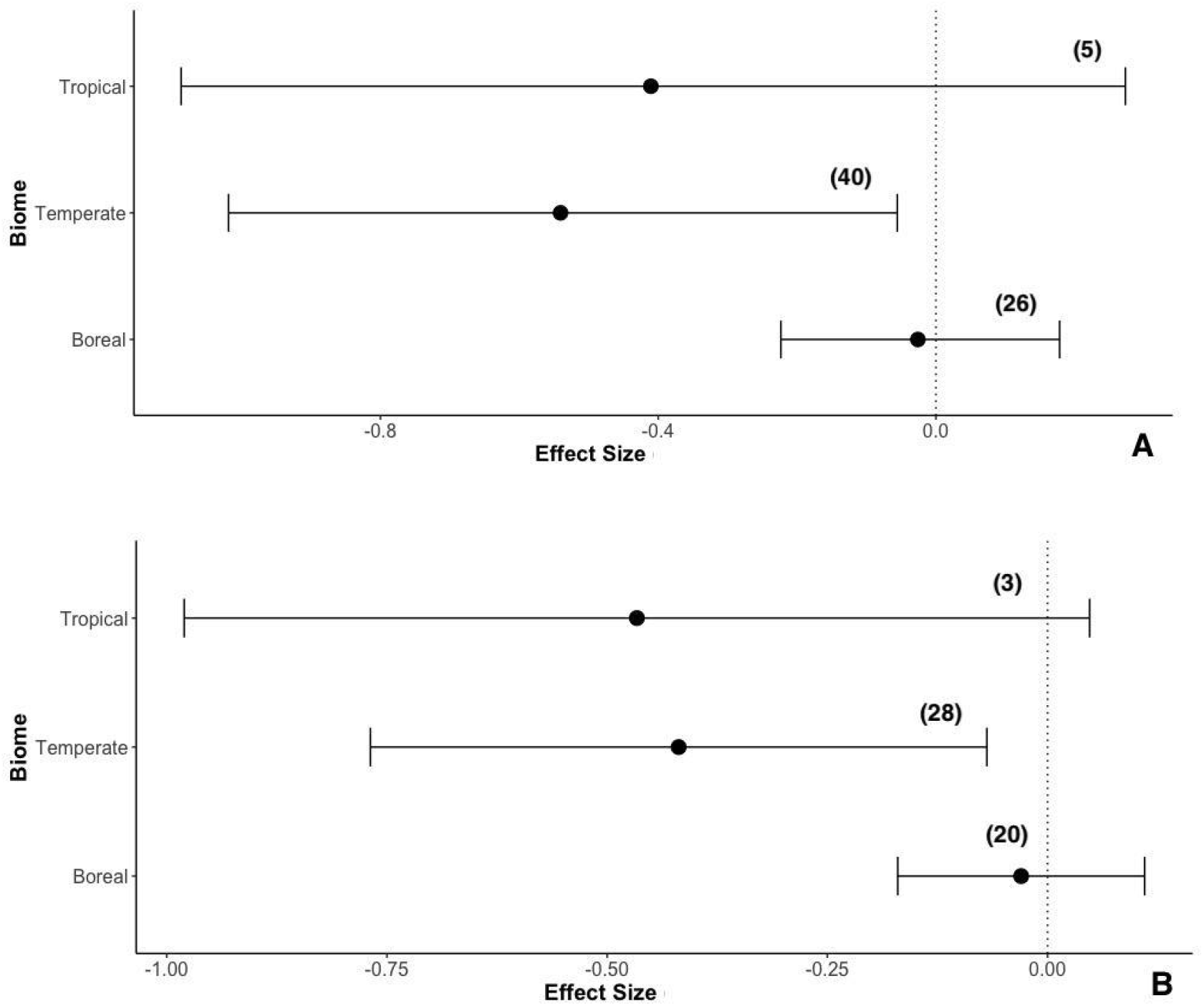
Effect of biome on effect size (Hedges’ *d* – mean effect size and 95% confidence intervals for each biome; tropical, temperate or boreal). In brackets is the number of studies in each group. A) Results for all studies reporting data on biome (total *k* = 71 studies). Biome was a significant predictor of effect size (Q_m_ for Biome = 13.17, *p* = 0.0014, *df* = 2). B) Results for low bias studies only (total *k* = 51 studies). Biome was a significant predictor of effect size (Q_m_ for Biome = 15.22, *p* = <0.001, *df* = 2).

None of the other moderators (Specialism, Life History or Alternative Hosts) had a significant influence on effect size (Figs 6-8). Damage by generalist pathogens was significantly lower in mixed forests compared to monocultures, but the effect was not significantly different from specialist pathogens (Fig. 6). Similarly, damage by biotrophic pathogens was significantly lower in mixed forests, but the effect was not significantly different from necrotrophic pathogens, which were not significantly affected by tree diversity (Fig. 8).

**Figure 6.**
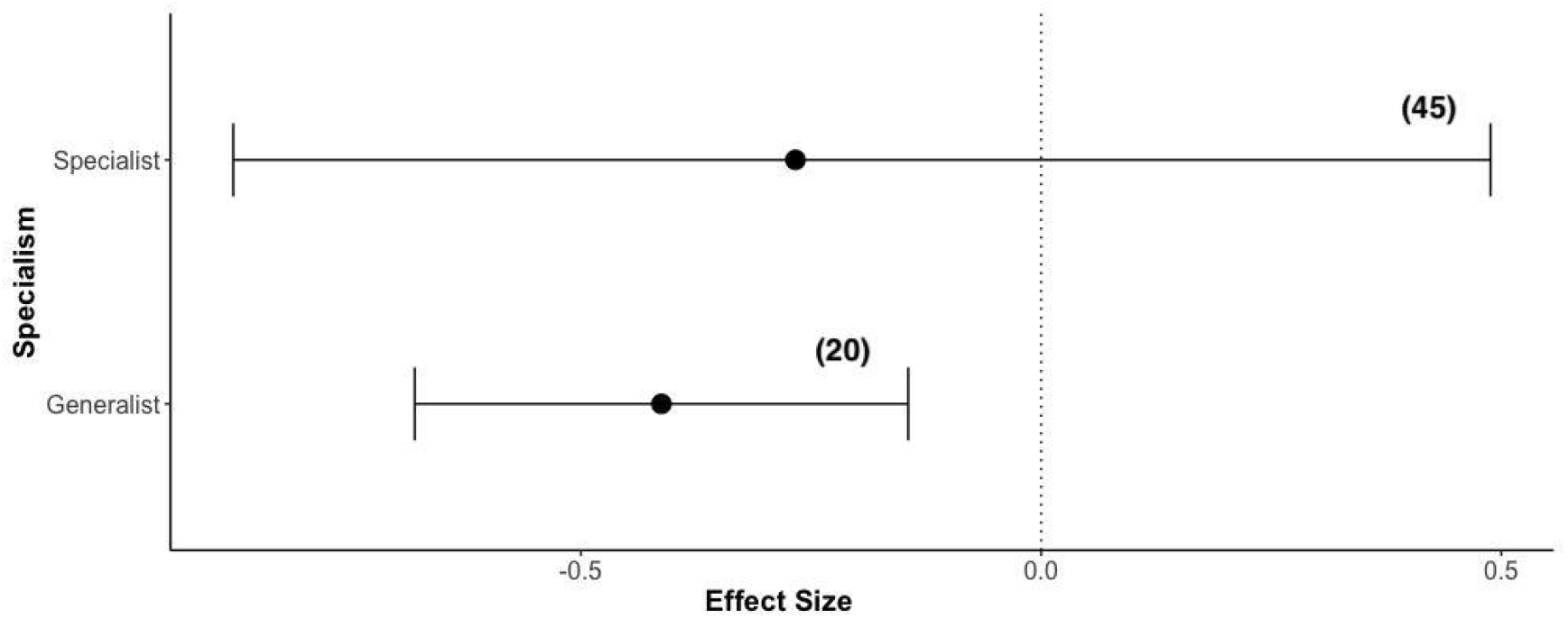
Effect of pathogen specialism on effect size (Hedges’ *d*). Shown is the mean effect size and 95% confidence intervals for pathogen specialists and generalists. In brackets is the number of studies in each group (total *k* = 65 studies). Pathogen specialism was a non-significant predictor of effect size (Q_M_ = 0.69, *p* = 0.41, *df* = 1).

**Figure 7.**
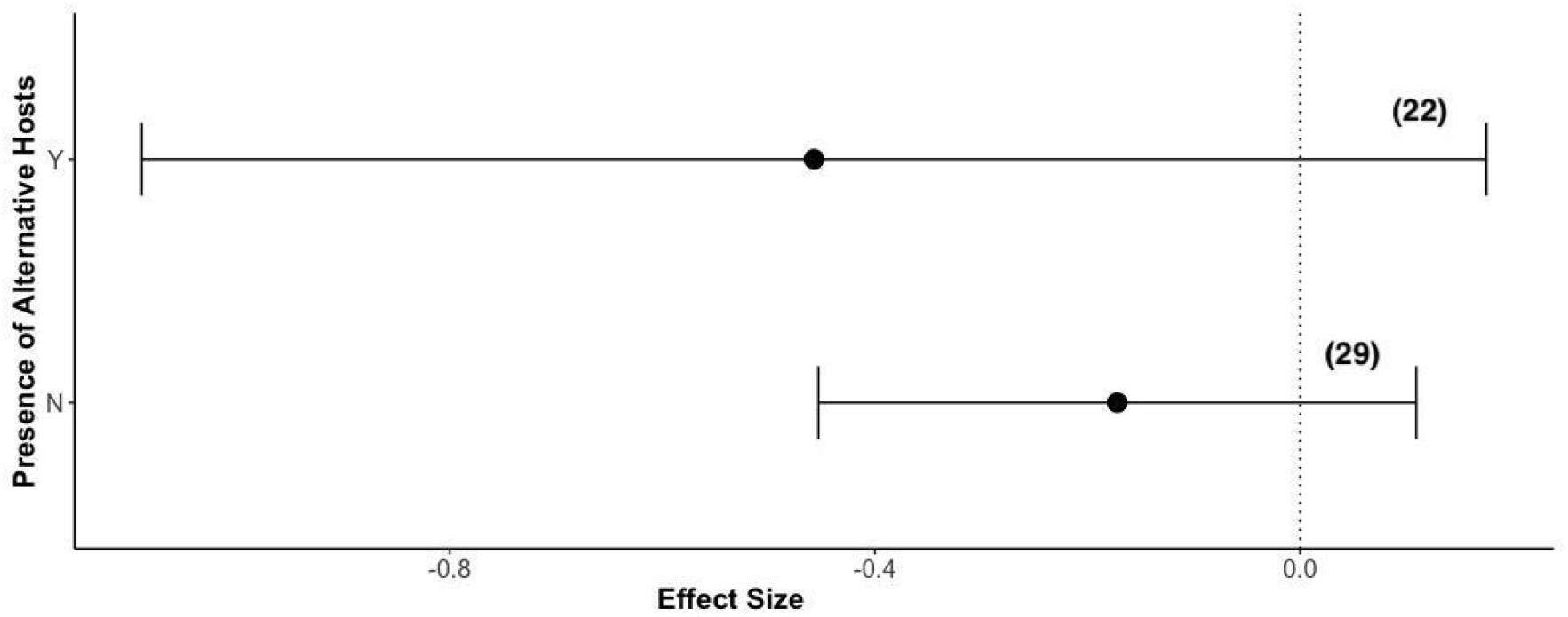
Effect of presence of alternative hosts (Y/N) on effect size (Hedges′ d). Shown is the mean effect size and 95% confidence intervals for each group. In brackets is the number of studies in each group (total *k* = 51 studies). Presence of alternative hosts was a non-significant predictor of effect size (Q_M_ = 2.53, *p* = 0.11, *df* = 1).

**Figure 8.**
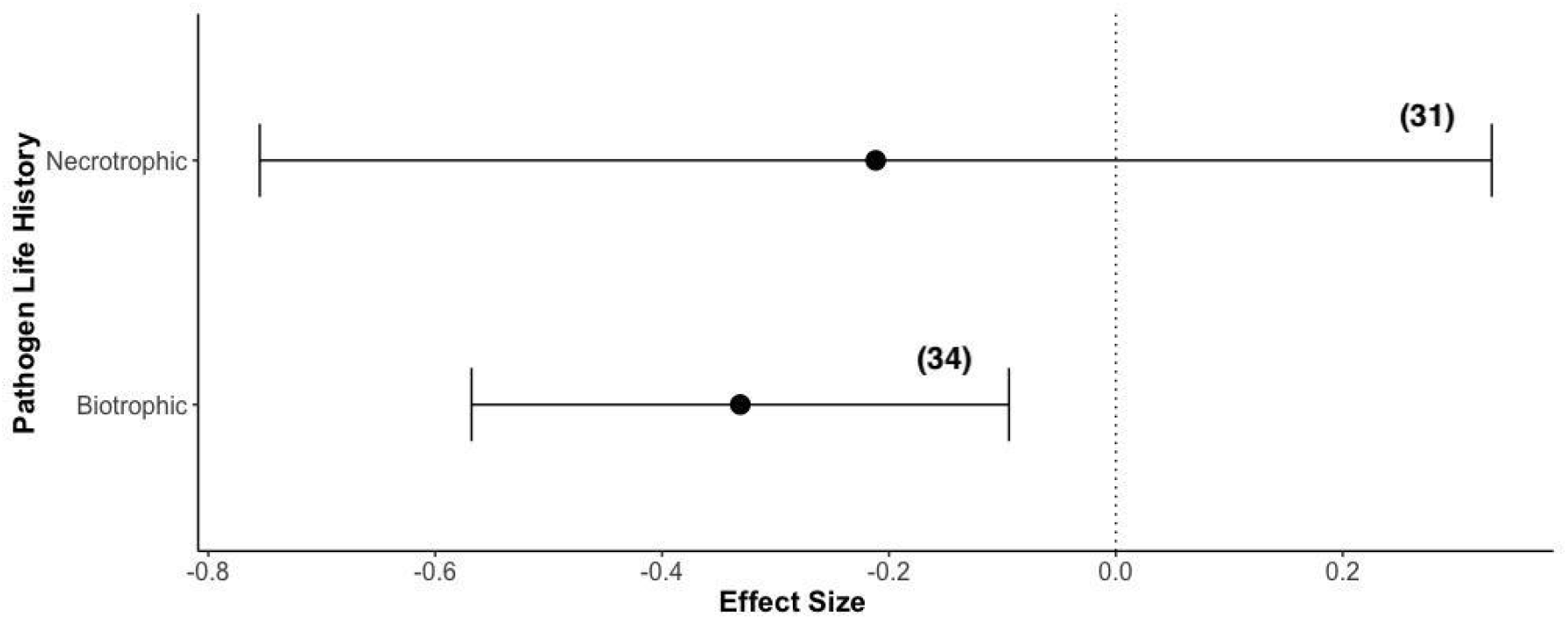
Effect of pathogen life history on effect size (Hedges’ *d*). Shown is the mean effect size and 95% confidence intervals for necrotrophs and biotrophs. In brackets is the number of studies in each group (total *k* = 65 studies). Pathogen life history was a non-significant predictor of effect size Q_M_ = 0.58, *p* = 0.45, *df* = 1).

Different levels of moderators were not significantly different from each other for any of the other variables that were extracted as metadata in our exploratory analysis (see Appendix V, Figs S2-S5).

### 3.3 Sensitivity analysis

Studies with a high versus low risk of bias did not differ significantly in their effect sizes (Fig. 9). However, studies with high risk of bias had significantly negative effect sizes whereas results from studies with low bias varied greatly. To test whether the main conclusions still held when high bias studies were removed, we re-ran the meta-analysis on low bias studies only. Removal of high bias studies removed the significant protective effect of mixed stands compared to monocultures (*d* = −0.17, 95% CI −0.39 to 0.048, *k* = 54 studies). But when low bias studies were again split into groups by biome (again removing studies occurring across biomes) there was a significant effect of mixed stands in temperate regions only, as the results of the main analysis had shown (Fig. 5B). Studies with converted/non-converted effects did not significantly differ in their effect sizes (Fig. S5).

**Figure 9.**
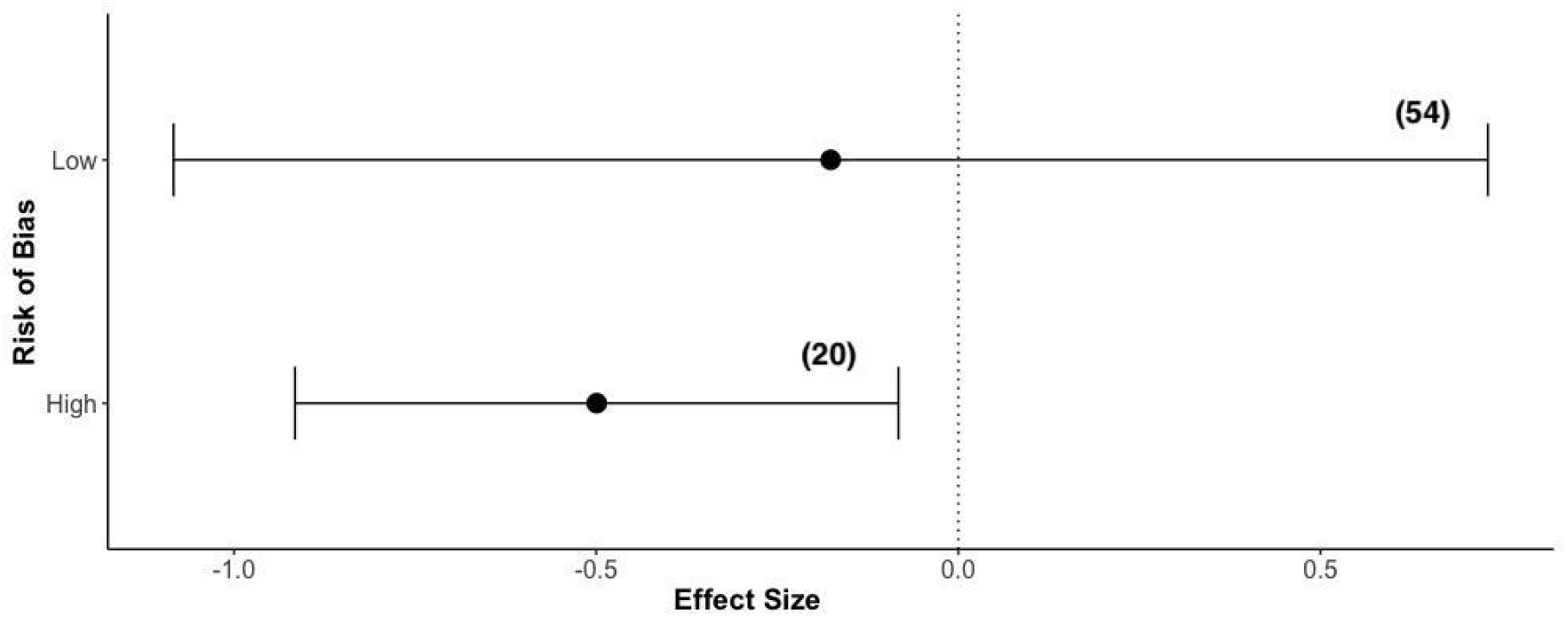
The effect of risk of bias on effect size. Shown is the mean effect size and 95% confidence intervals for low and high risk studies. Risk of bias as assessed *a priori* was a non-significant predictor of effect size (Q_M_ = 1.66, *p* = 0.20, *df =* 1).

### 3.4 Publication bias

Funnel plots suggested that there could be a negative correlation between effect size and standard error, as shown by the empty space in the bottom right side of the funnel plot of effect size against study standard error (Fig. S6). This space should have been populated by small, nonsignificant studies where effect size was positive but standard error large, i.e., studies that may have been subject to the “file-drawer” problem.

Egger’s regression test confirmed the significance of funnel plot asymmetry (*t* = -2.10, *p* = 0.039). This indicates that there may be some potential publication bias in this field. However, the overall mean effect (Hedges’ *d =* −0.27) was robust to publication bias, as shown by the high number of additional studies that would need to be added to render the overall effect size non-significant (Rosenthal’s fail safe *N =* 1310).

### 3.5 Qualitative evidence synthesis

Among qualitative studies, there were reports of both increases and decreases in pathogen damage in mixed stands. By their nature, qualitative studies did not contain sufficient quantitative data comparing mixed stands with monocultures to be included in a rigorous meta-analysis, so we did not attempt to analyse effect sizes or quantities of qualitative studies reporting positive or negative effects. Instead, we focused on extracting the suggested mechanisms of diversity effects reported in qualitative studies, to support future quantitative research.

Several qualitative studies were early field studies containing field reports (e.g. Day, 1927; Osmaston, 1927). Other qualitative studies were reviews of previous literature, such as reviews of long-term experiments or groups of experiments with the same theme (McCracken & Dawson, 1998; McCracken, Dawson, & Carlisle, 2005; Mundt, 2005). Some reviews cited other work, much of which was not available at full text in English when citations were searched (Korhonen et al., 1998; Pautasso et al., 2005). Most qualitative studies fell into groups of the same purported mechanisms to explain diversity effects on pathogens (Table 1).

**Table 1.**
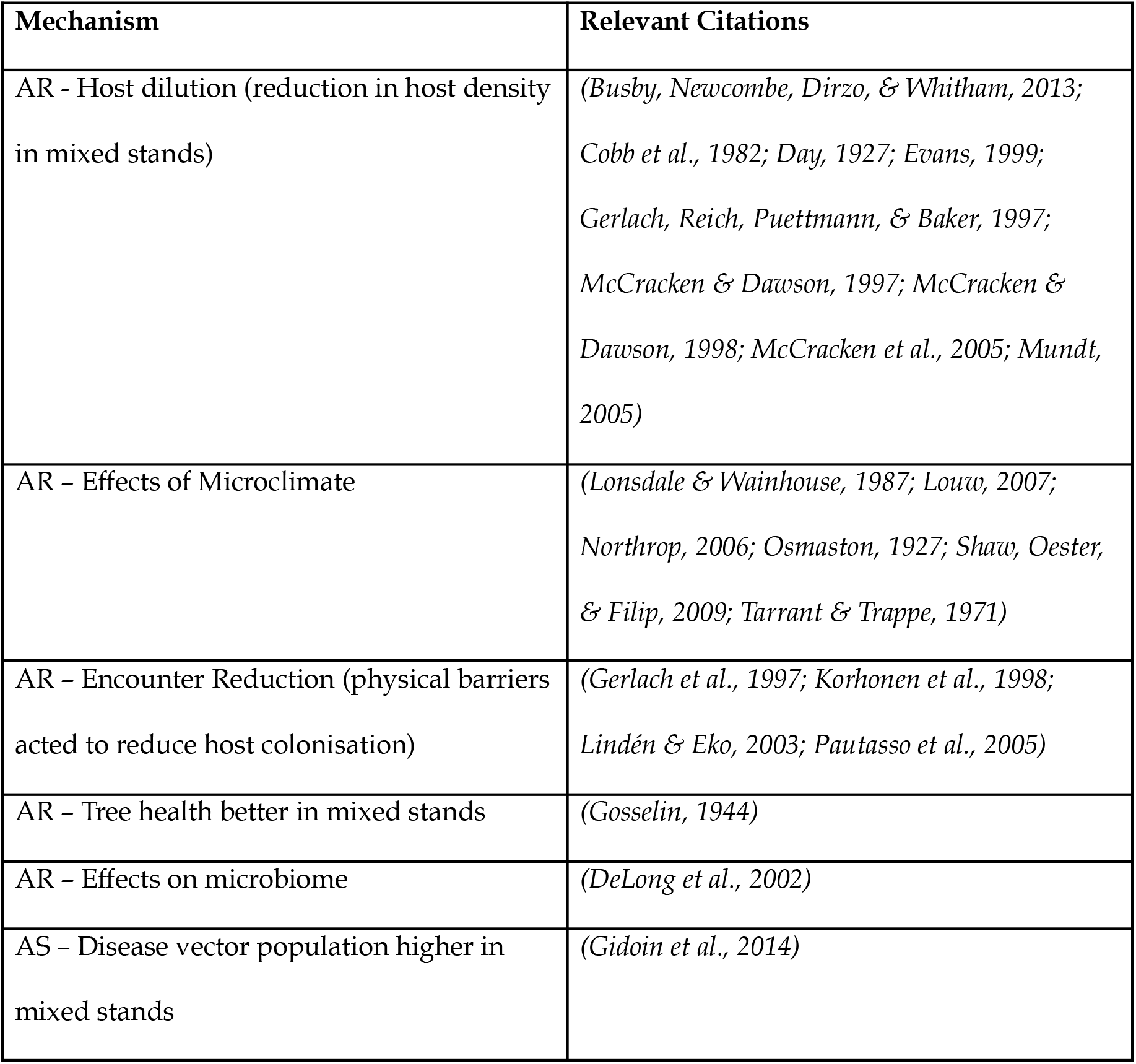
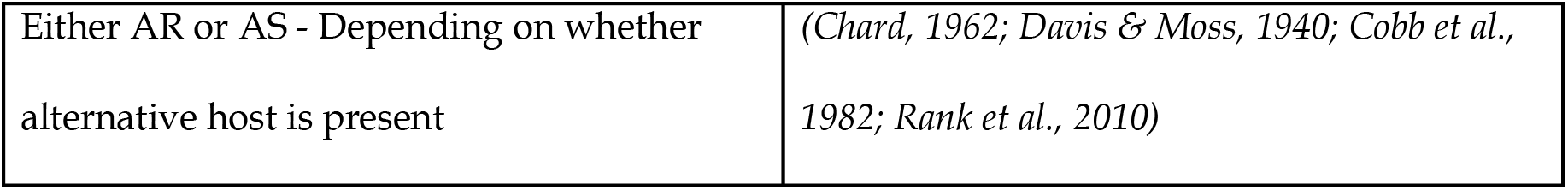
Mechanisms behind diversity-disease effects reported in qualitative papers, with citations provided for which that mechanism was reported. See discussion for description of the mechanisms, and the dataset associated with this paper for results and metadata of individual studies. AR = Associational resistance; AS = Associational susceptibility.

| **Mechanism** | **Relevant Citations** |
| --- | --- |
| AR - Host dilution (reduction in host density in mixed stands) | *(Busby, Newcombe, Dirzo, & Whitham, 2013; Cobb et al., 1982; Day, 1927; Evans, 1999; Gerlach, Reich, Puettmann, & Baker, 1997; McCracken & Dawson, 1997; McCracken & Dawson, 1998; McCracken et al., 2005; Mundt, 2005)* |
| AR – Effects of Microclimate | *(Lonsdale & Wainhouse, 1987; Louw, 2007; Northrop, 2006; Osmaston, 1927; Shaw, Oester, & Filip, 2009; Tarrant & Trappe, 1971)* |
| AR – Encounter Reduction (physical barriers acted to reduce host colonisation) | *(Gerlach et al., 1997; Korhonen et al., 1998; Lindén & Eko, 2003; Pautasso et al., 2005)* |
| AR – Tree health better in mixed stands | *(Gosselin, 1944)* |
| AR – Effects on microbiome | *(DeLong et al., 2002)* |
| AS – Disease vector population higher in mixed stands | *(Gidoïn et al., 2014)* |
| Either AR or AS - Depending on whether alternative host is present | (Chard, 1962; Davis & Moss, 1940; Cobb et al., 1982; Rank et al., 2010) |

## 4 Discussion

### 4.1 Mixed stands lower pathogen damage in temperate forests

Overall, we found a significant reduction in pathogen damage in mixed stands compared to monocultures, although the effect size varied from strongly negative to strongly positive (Fig 4). A significant negative effect size was only found in the temperate biome (Fig. 5), where 40 out of 74 studies were conducted (Fig. 2A-B). The reduction in pathogen damage in mixed stands was consistent in temperate forests, regardless of study bias (Fig. 5B). This suggests that the result for temperate biomes is the most robust pattern.

To our knowledge, this is the first meta-analysis providing evidence for a significant negative effect of tree diversity on pathogen damage in any biome. Liu et al. (2020), in a meta-analysis of plant diversity effects on infectious diseases, had previously reported a non-significant result in forests. They suggested that for forest trees, the identity of immediate neighbours is more important than overall plant community diversity in predicting disease damage, as several papers had noted the importance of tree neighbour identity effects on pathogens (Dillen et al., 2016; Hantsch et al., 2013).

However, their meta-analysis included estimates from only six papers on forests, all from higher latitudes (northern hemisphere). Their search was also confined to two databases for sources of studies (Web of Science and China National Knowledge Infrastructure).

The nonsignificant effect of tree species diversity on pathogen damage in the boreal biome contrasts to the findings of Nguyen et al., (2016) who found that tree species richness effects on pathogens were stronger at higher latitudes. Unfortunately, we could not test the effects of latitude or other climatic variables such as temperature or precipitation, as this would have required accurate latitudinal data to be available for all studies. Moreover, as the latitudinal gradient of studies was skewed towards the northern hemisphere, the gradient of precipitation and temperature that could have been tested would have been fairly narrow. Interestingly, although fewer studies were conducted in boreal forests than temperate forests, they showed less variance than both tropical and temperate forests (Fig. 5). This suggests that the nonsignificant result for boreal versus temperate forests is likely to be reliable (Fig. 5).

### 4.2 Pathogen specialisation is not a key driver of mixed stand effects for forest pathogens

Previous meta-analyses found that the effect of tree diversity on insects was stronger for specialists compared to generalist insect herbivores (Castagneyrol et al., 2014; Jactel & Brockerhoff, 2007). This supports the hypothesis that dilution effects are stronger when there is a lower likelihood of alternative hosts for a pest being present in a stand. In contrast, we did not find evidence that the same trend held true for forest pathogens.

Neither pathogen specialism nor the presence of alternative hosts in stands had significant effects on pathogen damage in mixed stands compared to monocultures. This suggests that the negative tree diversity effect on pathogens is not purely caused by a dilution effect. However, considering generalist pathogens on their own, tree species diversity did have a significant negative effect (Fig. 6).

This initially surprising result for generalist pathogens could have several explanations. The majority of studies of generalist pathogens (16 out of 20 studies) were of two important groups of forest pathogens. These were *Phytophthora ramorum* (3 studies), and the root rot pathogens *Armillaria* sp. and *Heterobasidion annosum* (5 and 8 studies respectively) that affect multiple conifer and broadleaved species. The negative effect of tree diversity on *Phytophthora ramorum* damage to host species was documented in landscape-scale studies from California, USA (Dillon & Meentemeyer, 2019; Haas et al., 2016; Haas, et al., 2011). Previous work had shown that although *P. ramorum* is a generalist pathogen, host damage is predicted by the density of its most competent host (bay laurel) in stands (Cobb et al., 2010; Rank et al., 2010). Thus, at the landscape level, the studies included in the meta-analysis here found a reduction in pathogen damage at greater tree diversity, as bay laurel density reduced (Dillon & Meentemeyer, 2019; Haas et al., 2016; Haas et al., 2011). As many tree diversity experiments are planted in plots of limited size, it has been suggested that edge effects between treatments or the surrounding landscape could be eroding the effects of tree diversity (McCracken et al., 2005). To predict the effects of tree species diversity on pathogen damage in a particular forest or stand, it may be important to take into account epidemiological factors at a landscape-level, such as the presence of alternative hosts nearby.

The negative impact of tree diversity on root rot pathogens, which impact numerous coniferous species worldwide, has also been well documented in previous reviews, although occasional increases in damage have also been observed (Korhonen et al., 1998; Pautasso et al., 2005). One hypothesis is that there is a reduction in disease transmissibility due to encounter reduction in the presence of alternative species, as root rot pathogens are spread via root-to-root contacts between susceptible hosts (Korhonen et al., 1998; Pautasso et al., 2005, Table 1). Several qualitative studies of root rot pathogens suggested the importance of considering tree neighbour identity. These studies suggest that increasing proportions of non-host broadleaved species in stands can have a stronger disease reduction effect (Lindén & Eko, 2003; Pautasso et al., 2005).

However, 2 out of 3 studies of root rot pathogens with positive effect sizes in the meta-analysis had coniferous species planted in mixtures, rather than broadleaves, suggesting this is not a universal mechanism to explain associational resistance (Arhipova et al., 2011; Korhonen et al., 1992; Whitney, 1973). Moreover, in the meta-analysis, the presence of alternative hosts in stands did not significantly influence effect size. Although the presence of alternative hosts may reduce the impact of tree diversity in individual cases, it is not a general trend that can be relied upon for predictive forestry.

Rather than the presence of alternative hosts, other direct or indirect effects of tree neighbour identity, such as their effect on the soil microbiome, or microclimatic conditions, may underlie their associational effect on forest pathogens. For instance, Tarrant & Trappe (1971) suggested that the presence of alder in mixed stands could reduce damage by the root rot pathogen *Phellinus weirii* on *Pseudotsuga menziesii* and other conifers, by producing phenolics and other ammonia-containing compounds to suppress fungal growth. In another example, the presence of paper birch in mixed stands with Douglas Fir has long been documented to reduce *Armillaria* damage to Douglas Fir (Baleshta et al., 2005; Louw, 2007). DeLong et al., (2002) found that *Armillaria-*inhibiting populations of pseudomonad bacteria increased around birch roots, suggesting that paper birch was altering the microbiome to create a “suppressive soil” for *Armillaria.* In a case of associational susceptibility in mixed stands, the authors suggested that the presence of hemlock in mixture with beech increased the humidity of stands and protected beech boles from winter freezing, both of which would benefit the scale insect vector of beech bark disease (Twery & Patterson, 1983). A previous study by the authors (Field et al., 2020) also found that the species composition of mixed stands altered the stand microclimate (temperature and humidity) and was correlated with oak powdery mildew infection (a more stable microclimate increased mildew damage). The environmental conditions created in mixed stands, dependent on the identity of neighbouring tree species, may therefore be as important as their potential as alternative hosts for disease.

Where multiple hosts are required to complete the pathogen life cycle, some qualitative studies indicated that the impact of mixed stands depended upon whether alternate hosts were planted. For instance, rust species such as white pine blister rust (*Cronartium ribicola)* require both *Pinus strobus* (the aecial host) and *Ribes* sp. (the telial host) to complete their life cycle. In the first half of the twentieth century in the United States and United Kingdom, large-scale removal of naturally regenerating *Ribes* sp was one of the few known treatments for white pine blister rust, but was eventually abandoned as an approach for economic reasons (Chard, 1962; Davis et al., 1940). In this case, planting white pine as a monoculture is protective, as the disease cannot complete its life cycle without the presence of the alternative host. The same is true for pine twist rust (*Melampsoa pinitorqua*) for which the other main host is aspen. A study from Finland therefore found when *Pinus strobus* was planted in mixture with aspen, associational susceptibility was the result compared to monoculture (Mattila, 2005).

### 4.3 Biotrophic pathogens are reduced in mixed stands

We found that biotrophic pathogens caused significantly lower damage in mixed stands, whilst necrotrophic pathogens were not significantly affected. While necrotrophic pathogens varied, studies of biotrophic pathogens (that rely on a living host to complete their life cycle) were exclusively of powdery mildew and rust species (34 out of 34 studies). This group of pathogens has been extensively studied in forest, crop and grassland systems (Mundt, 2005), perhaps as they are easy to identify and score in the field with strong visual symptoms. As rust and powdery mildew species are passively wind-dispersed species, epidemiological theory would suggest that the success of dispersal will depend upon the distance between susceptible hosts, although this theory has yet to be tested in the forest environment (Mundt, 2005).

### 4.4 Host, pathogen, and environment

The disease triangle, a key concept in plant pathology, describes the outcome of a disease to be dependent upon triangular interactions between host, pathogen and environment (Francl, 2001; Stevens, 1960). Assimilation of the evidence on mixed stands and forest pathogens reveals the sheer variety of outcomes that are possible when increasing tree diversity, and the many potential mechanisms underpinning such effects on pathogens. We suggest the use of the disease triangle as an aid for ecologists and foresters who wish to conduct hypothesis-focused research. Figure 10 shows how such research could begin by considering the impacts of mixed stands on these three areas for any given pathosystem. Host, pathogen and environment may all be impacted in different ways by the planting of species mixtures, with the overall outcome requiring assessment of all three areas.

**Figure 10:**
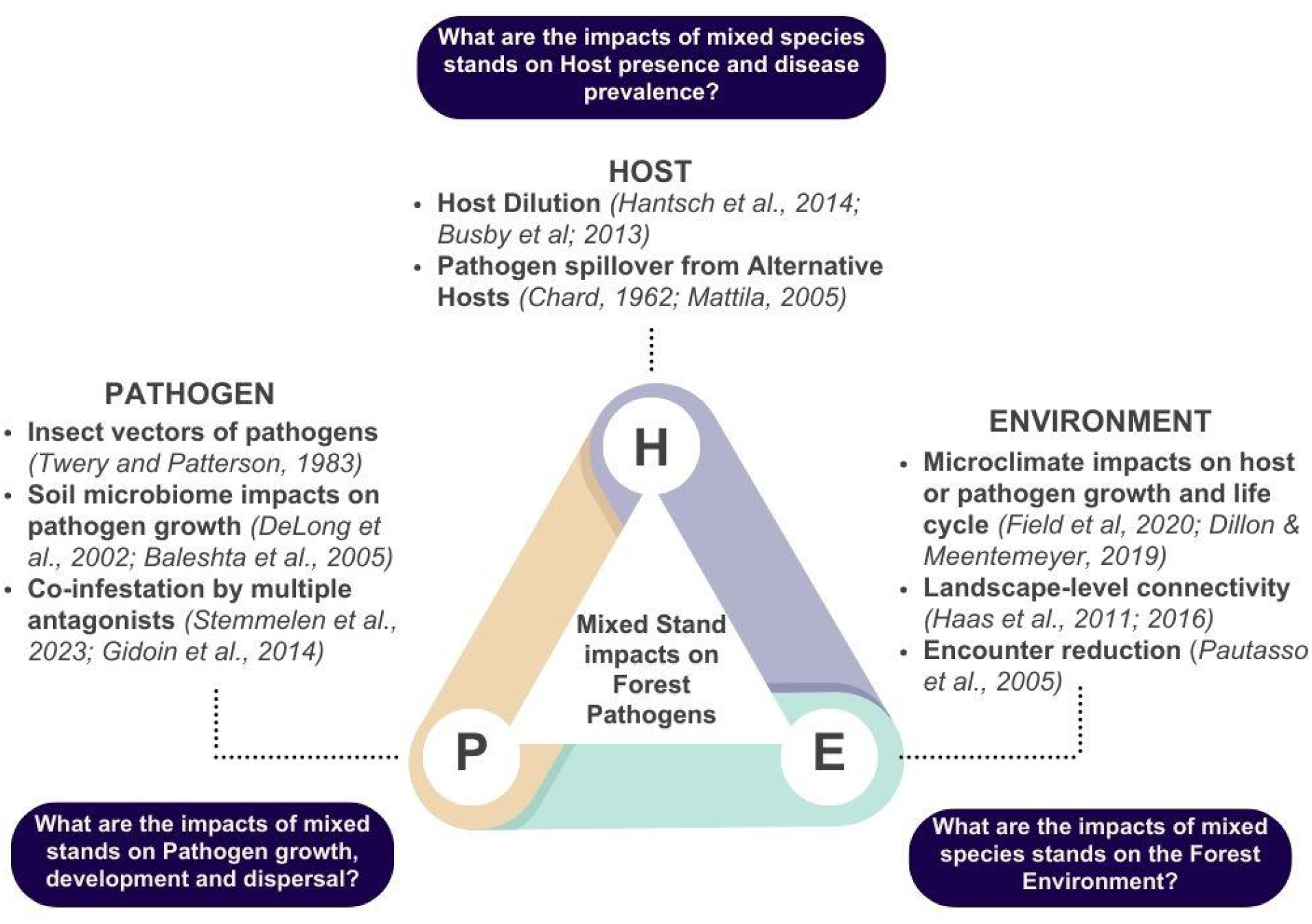
Summary of factors to consider in the outcome of mixed stands on forest pathogens, arranged around the classical disease triangle of pathogen, host and environment. Citations are example papers which illustrate a mechanism.

### 4.5 Knowledge gaps

There was an obvious latitudinal bias present in our studies, with just 5 out of 74 studies in the meta-analysis from the tropical biome and 4 out of 30 qualitative studies from the tropics. There was no significant effect of mixed stands on pathogens in tropical biomes, but the sample size was perhaps too small to ascertain if this was a real trend with only 5 studies from the tropics (Fig. 5). In contrast to our results, Liu et al. (2020) had suggested weaker plant diversity effects on pathogens with increasing latitude, although their study did not contain any tropical studies. They had postulated that greater competition due to higher overall plant diversity at lower latitudes could have led to higher host specialisation and therefore strong effects of resource dilution (Liu et al., 2020). Our results find no evidence to support this conclusion, although the lack of accurate latitudinal data for all studies made it difficult to disentangle the mechanisms behind biome effects. More research on the effect of mixtures on disease in tropical plantations could have a beneficial impact for small-scale farmers. For instance, Dumont et al., (2014) found in a social study of cocoa farmers that 2% of farmers reported an increase in black pod disease with increasing tree diversity, while 6.8% reported decreases, showing that a large proportion of farmers remained neutral or lacked sufficient knowledge to make planting decisions.

Unsurprisingly, studies were also biased almost entirely towards fungi, which are the most widespread and economically important group of plant pathogens on the planet (Fisher et al., 2012). There were only a small number of studies of *Phytophthora* (oomycete) species (7 studies total), despite this genus including some of the most lethal group of plant pathogens in the world (Brasier, 2008). The one study of bacteria included in the meta-analysis was also one of only two quantitative studies from South America (Chile), reporting a negative but nonsignificant effect of tree diversity on *Pseudomonas siringae pv. Mors-prunorum* damage to *Prunus avium* (Loewe, González, & Balzarini, 2013). There is therefore clear potential for future research on the effects of mixed stands on bacterial and oomycete species.

In our results, there was a notable lack of studies with small, nonsignificant positive effect sizes (Fig. S6). This is consistent with the predictions of a “file drawer problem” – that small, nonsignificant studies are rarely published due to institutional and journal bias against such studies (Rosenthal, 1979). Previous reviews in this area have consistently discussed the benefits of mixed stands on pest and pathogen damage (e.g. Jactel et al., 2017; Roberts et al., 2020). While this may be a real trend, the fact that we found a bias against studies with large standard errors suggests that some publication bias could be at play. A possible consequence of this could be an overestimation of the true effect in the literature.

### 4.6 Study limitations and suggestions for future work

In our study, we could not directly test whether one of the major mechanisms suggested by authors of previous studies held true: that associational resistance was driven by a reduction of host density (resource dilution) in mixed stands. This was due to host density often not being reported in studies, although we may be able to assume that host dilution often increased with increasing tree diversity. However, the fact that both pathogen specialism and presence of alternative hosts had nonsignificant impacts on the effect size, and that a significant effect of mixed stands was still seen for generalist pathogens, suggests that the effect of mixed stands on forest pathogens is not always as simple as a host dilution effect even if host density is lower at greater diversity levels.

Instead, we found several studies suggesting the importance of tree neighbour identity for influencing pathogen damage on focal hosts. In fact, one study actually reported an increase in disease with a reduction in host abundance. The main vector of black pod disease (*Phytophthora megakarya*) of cacao trees (*Theobroma cacao*) are ants, which were found in lower numbers with decreasing plant diversity, therefore potentially lowering disease in monocultures (Gidoin et al., 2014). This also highlights the importance of considering how a disease is vectored, a step which is not so common for insects but that may include other trophic levels for forest pathogens.

Another major limitation of our study was the inability to mine data from papers that were not available in English. Several studies relevant at abstract level were not available at full text in English, in particular, many forestry studies from Eastern Europe, cited in reviews such as Korhonen *et al* (1998). These studies would almost certainly have been a source of additional data for meta-analysis. Forestry researchers should note the large quantity of studies from Eastern Europe that are not available for English systematic reviews and meta-analyses, representing a “black hole” of literature as yet unincorporated into the English-speaking areas of the scientific world. A future collaborative study between several multi-lingual scientists, with the aim of including this literature, could be highly valuable. For instance, a previous study of the invasive pest literature highlighted the importance of data mining literature from Chinese to English, when modelling the spread of insect pests in China (Bebber et al., 2019).

Finally, due to limitations of time and resources, this study was conducted with a dataset up to 2019 only. It is possible that in the intervening years, additional data has been published which could impact the size or direction of effect sizes measured.

However, the dates of studies reviewed here range from 1927 - 2019 and we underwent a rigorous systematic review process to find all relevant literature to that date. We are therefore confident that this review synthesises our understanding of the tree diversity-pathogen literature up to the very most recent years, providing valuable insights for practitioners and researchers alike.

## Conclusion

It is often assumed that, as is the case for insect pests, increasing tree diversity in forest plantations has a negative effect on forest pathogens. Using meta-analysis, we found a small but significant negative effect of mixed stands on tree disease, that was especially clear in temperate forests for which the most data was available. For some pathogens (for instance, *Armillaria* sp.*, H. annosum,* rusts and mildews), there was clear and plentiful evidence that increasing tree diversity reduced pathogen damage. The importance of tree neighbour identity rather than host dilution on its own became clear in our synthesis of both the quantitative and qualitative evidence. In the planting of mixed stands, it is therefore important to understand specific aspects of the study system at hand, rather than a general recommendation to plant mixed stands. Future experimental work in mixed forests could focus on unravelling associational effects by considering neighbouring tree species impacts on aspects such as forest microclimate and soil microbiome.

## Author contributions

EF & JK conceived the study and the methodology. EF carried out the data searches, data extraction and evidence synthesis. EF carried out the meta-analysis with input from JK and AH. EF analysed the results and wrote the manuscript with significant input from NB. All authors commented on the final version of the manuscript.

## Acknowledgments

We thank Hervé Jactel and colleagues who provided a list of relevant studies that helped develop the scoping searches. Thank you to all the authors who provided additional data from previously published studies for the meta-analysis: Pilar Fernandez-Conradi, Mike Cruickshank, Whalen Dillon, Diem Nguyen, Sarah Haas and Ross Meentemeyer. We also thank additional researchers from TreeDivNet for providing additional unpublished data: Matthias Dillen and Martin Schädler. This systematic review would not have been possible without the support of EF’s lab group, the Hector-Turnbull Plant Ecology lab at Oxford. Emma Jardine, Emily Warner, and April Burt assisted in the full text and abstract screening stages as additional reviewers for kappa tests. We are also grateful to Gill Petrokofsky for her advice and support in the development of systematic review methodology and to Neal Haddaway and independent peer-reviewers from the Centre for Environmental Evidence who reviewed the systematic review protocol. EF would like to thank her other PhD supervisors, Melanie Gibbs and Karsten Schönrogge for commenting on earlier versions of this manuscript. A big thank you to Forest Research UK for providing the additional funding needed to bring this manuscript to publication.

## Supplementary Material for Field, *et al*., (2024)

### Appendix I: Description of scoping searches and main literature search

A scoping study conducted using the database Scopus was used to identify the search terms for use in bibliographic databases and internet searches. This was done by testing search strings in Scopus using the Field Tag (TITLE-ABS-KEY) to search for relevant titles, abstracts and keywords. Search terms were sourced from previous meta-analyses and systematic reviews on themes of mixed forests and/or insect herbivory and forest pathogens, and from the keywords of studies identified *a priori* as highly relevant (see Appendix II). Search results were ordered based on relevance and limited to studies in the ‘Agriculture and Biological sciences’ and ‘Environmental Sciences’ topics to remove results from non-relevant subjects. For each search combination, the total number of hits and the number of relevant papers (assessed based on title) in the first 200 hits was recorded. In addition, each search combination was scrutinised for its ability to retrieve a list of **12** studies identified a priori as highly relevant to the primary question (test library, see below). These studies were sourced from the personal libraries of the authors and stakeholders. Although all contain information relevant to the primary question, they utilise a range of observational and experimental study systems and the phrasing related to the primary question differs in each case; not all combinations of search terms were able to retrieve all of these papers. The final search terms were able to retrieve all of these papers, and achieved a good balance between relevant and non-relevant studies in the first 200 results.

The final search terms and their relevance to each of the PECO question elements is summarised in Table S1. Table S2 shows the exact search terms used in each database, the number of hits returned when initially searched, and search date. The large discrepancy in numbers of hits returned from Scopus compared to Web of Science and CABI is likely to be due to the restricted field tag used (TITLE-ABS-KEY) combined with differences in the literature available from Scopus, as without use of this field tag the number of hits returned by this database are typically extremely high (a comparison search using the same search terms without this field tag in Scopus returned over 37,000 hits).

Due to the large number of search results from irrelevant topics, the results of the Scopus search was limited to journals in the “Agricultural and Biological Sciences” and “Environmental Science” categories. In Web of Science, results of searches were limited to the following research areas in order to narrow the search: Agriculture OR Forestry OR Environmental Sciences OR Ecology OR Plant sciences.

**Table S1:**
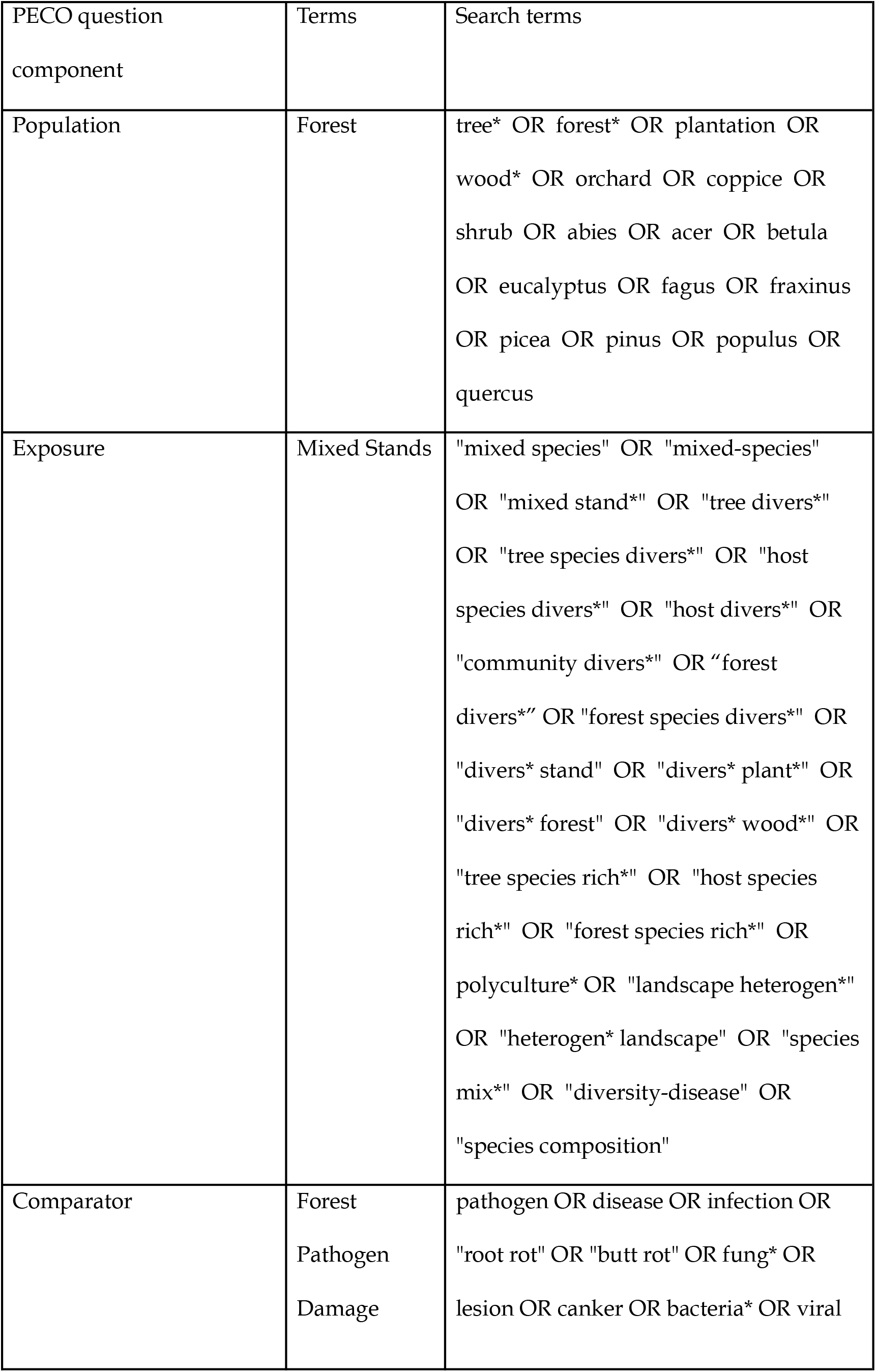
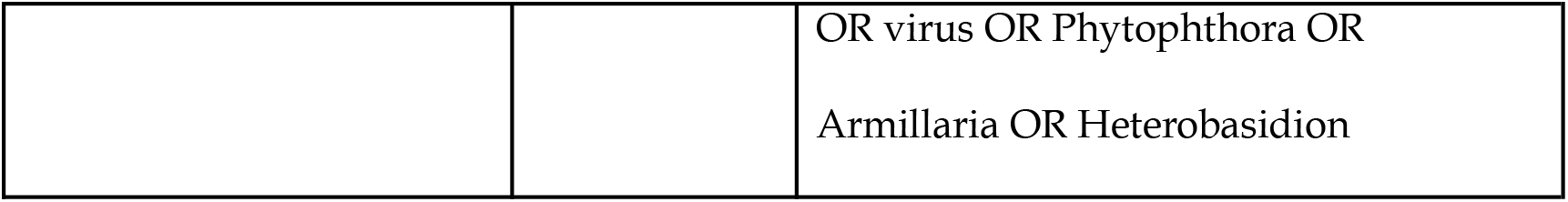
List of search strings used in bibliographic databases and their relationship to each component of the primary question. Each section was combined using ‘AND’ (for exact search strings with Boolean operators used in each database, see Table S2).

**Table S2:**
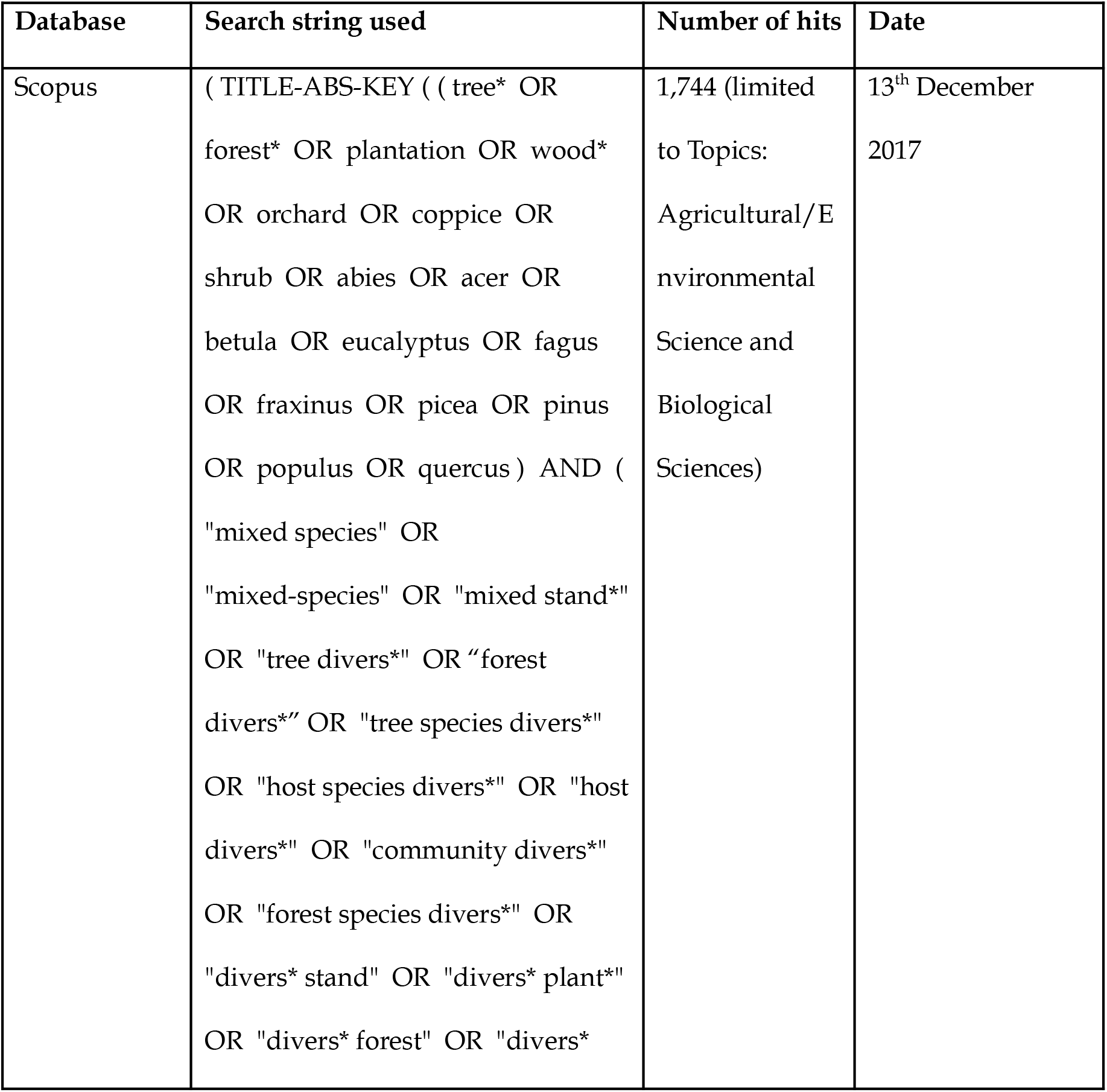
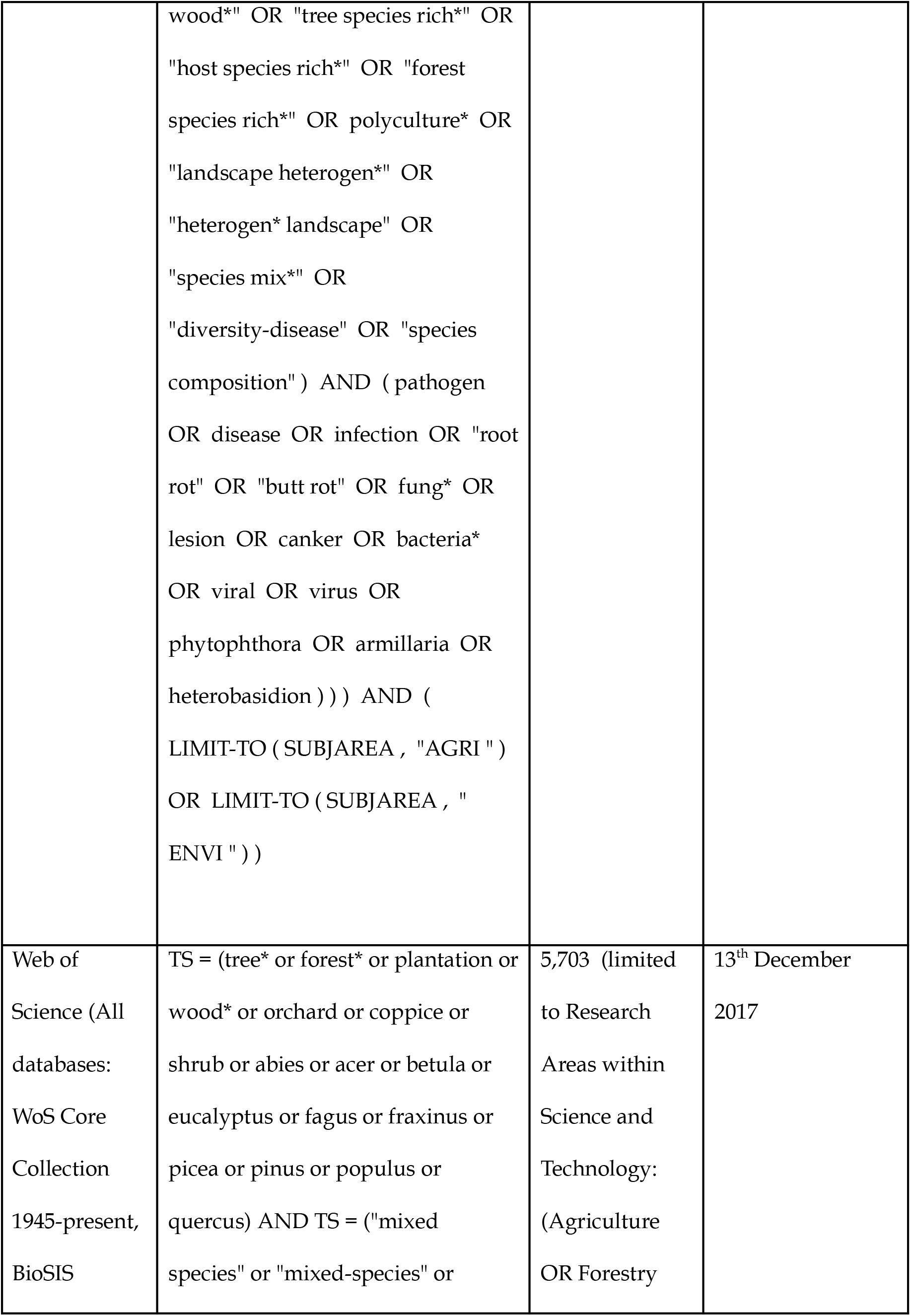
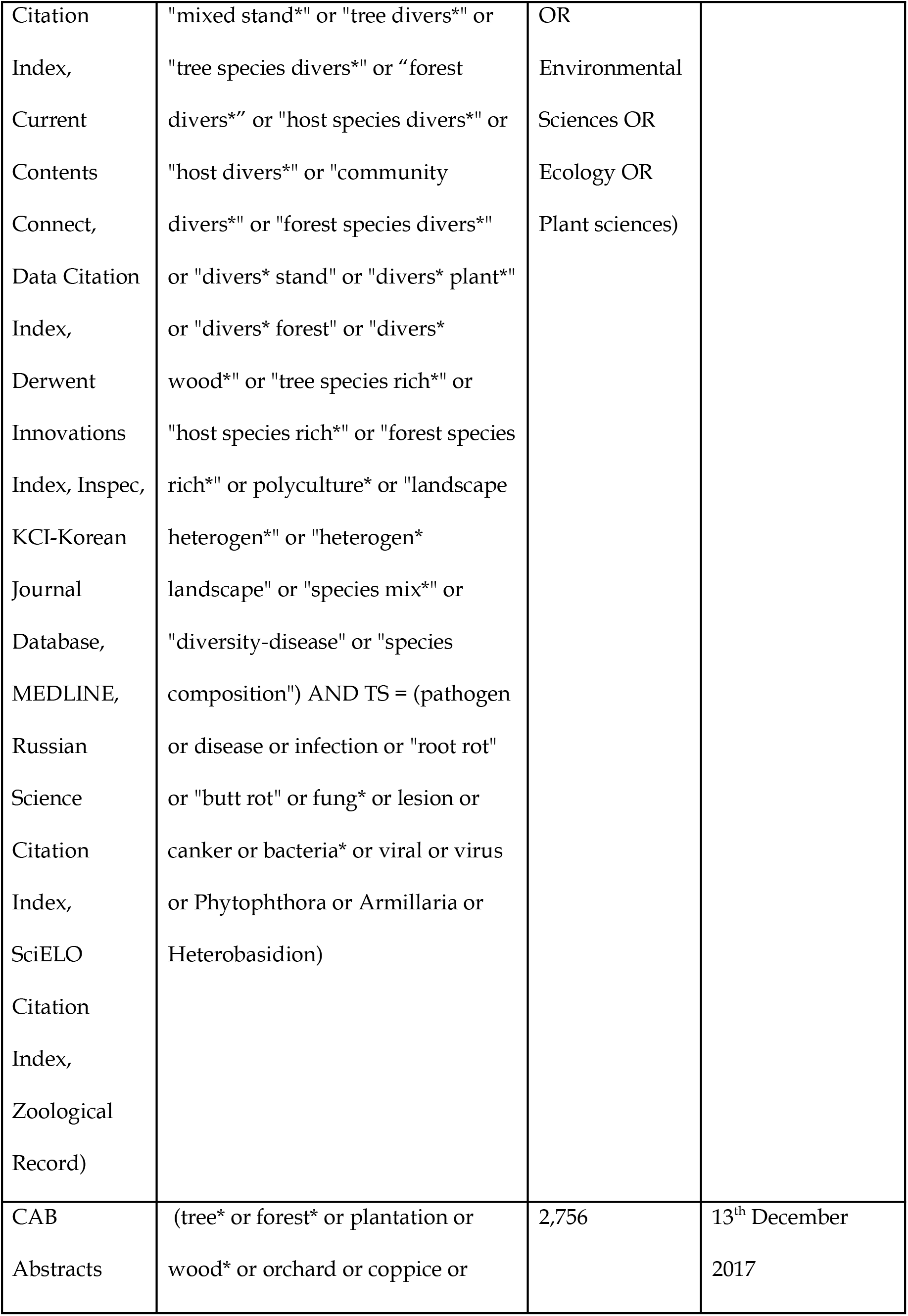
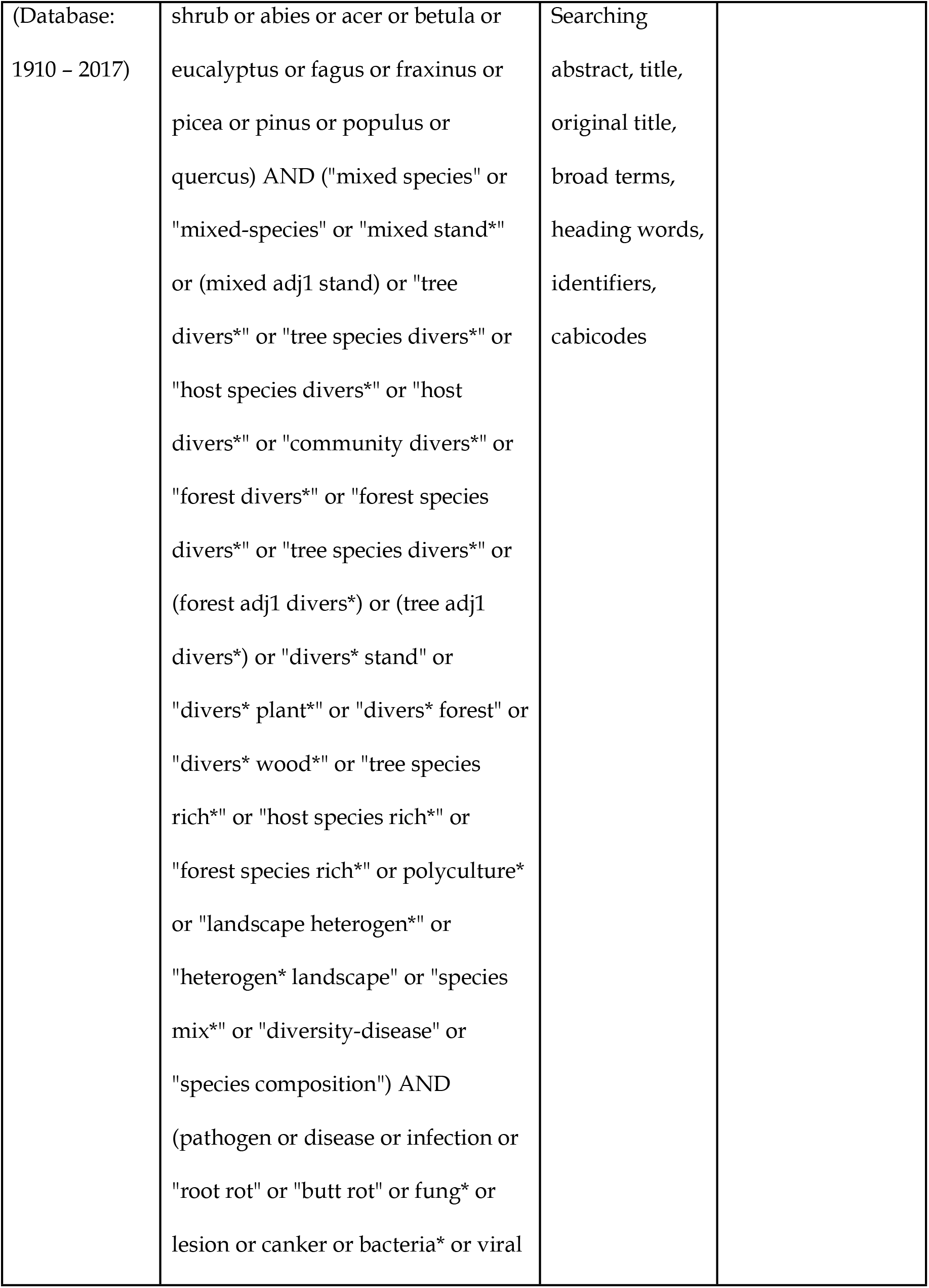
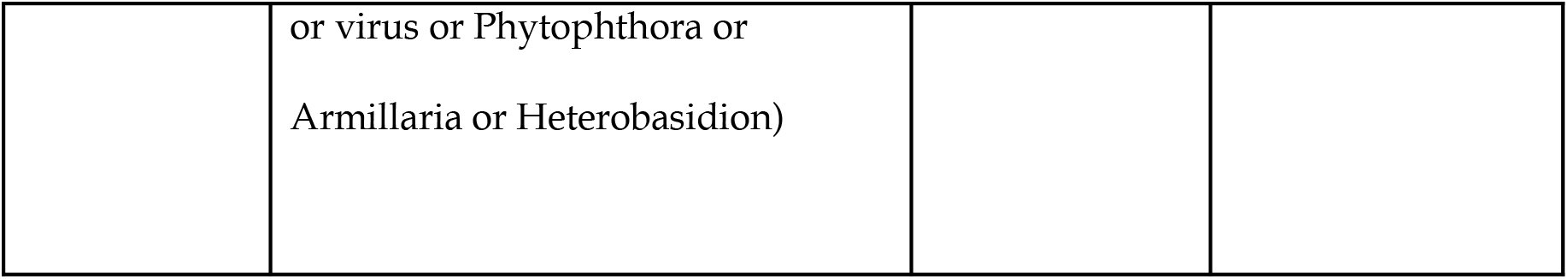
List of search strings used in each major database queried during the systematic review, with number of hits returned and date initially searched. NB: Google Scholar does not permit the use of Boolean operators, so search terms were modified and the first 1000 results downloaded from Google Scholar. We tried several search term combinations in Google Scholar to maximise the number of test library studies found while minimising duplication with the rest of the literature searches. Final search terms used in Google Scholar were: ‘ “mixed stand” forest pathogen ‘ (Search completed on 21^st^ February 2018).

### Appendix III: List of stakeholders consulted during the literature searching stage

Centre for International Forest Research (CIFOR)

EU COST action on Mixed Forests (EUMIXFOR)

EUFORGEN

European Forest Institute

Food and Agriculture Organisation of the United Nations Forest Europe

Forest Research (UK)

French National Institute for Agricultural Research (INRA)

International Union of Forestry Research Organisations (IUFRO)

United States Department of Agriculture

World Agroforestry Centre

### Appendix IV: Metadata Form sent to stakeholders from TreeDivNet contributing unpublished data sets

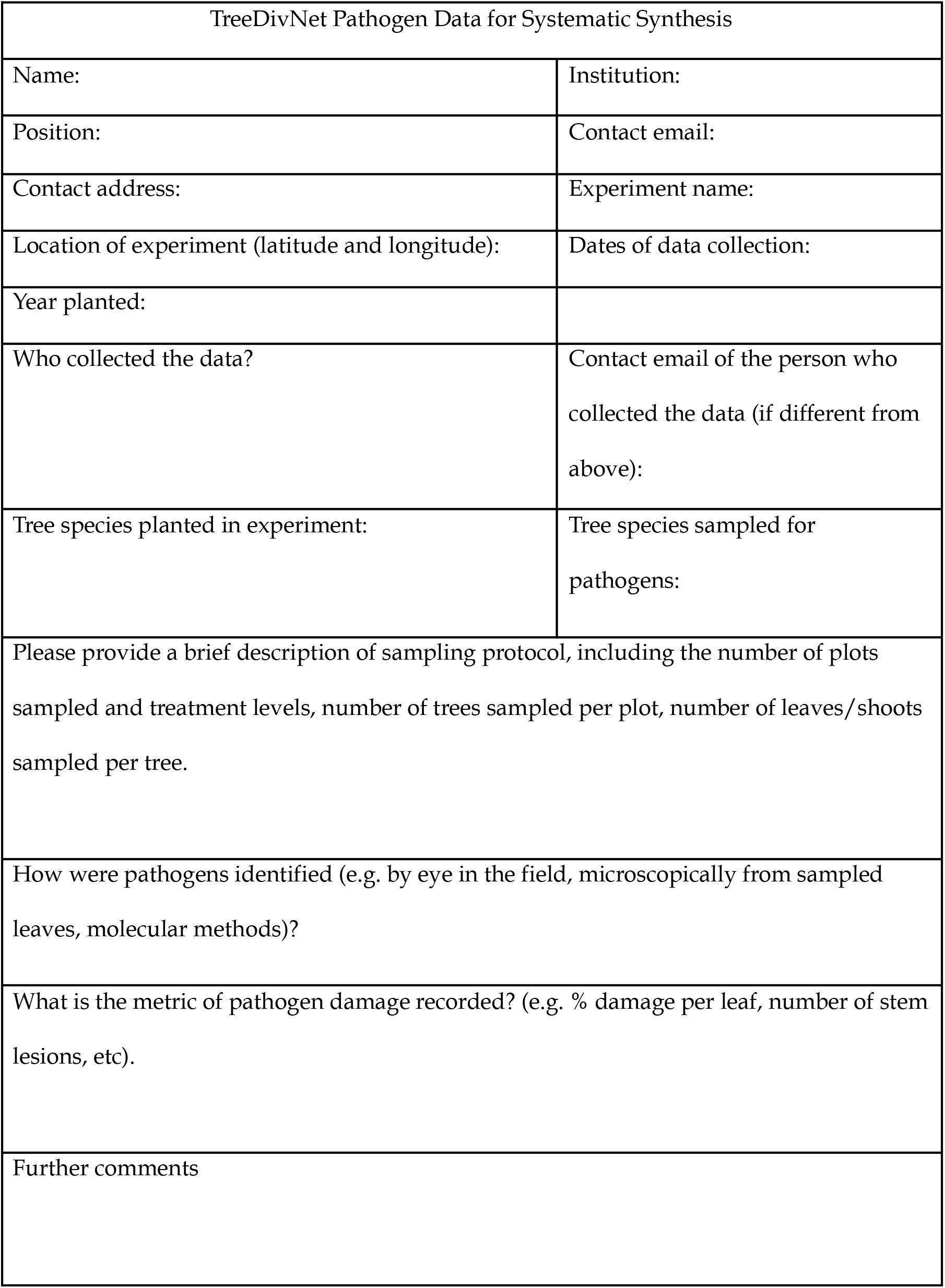

### Appendix V: Supplementary Figures

**Figure S1.**
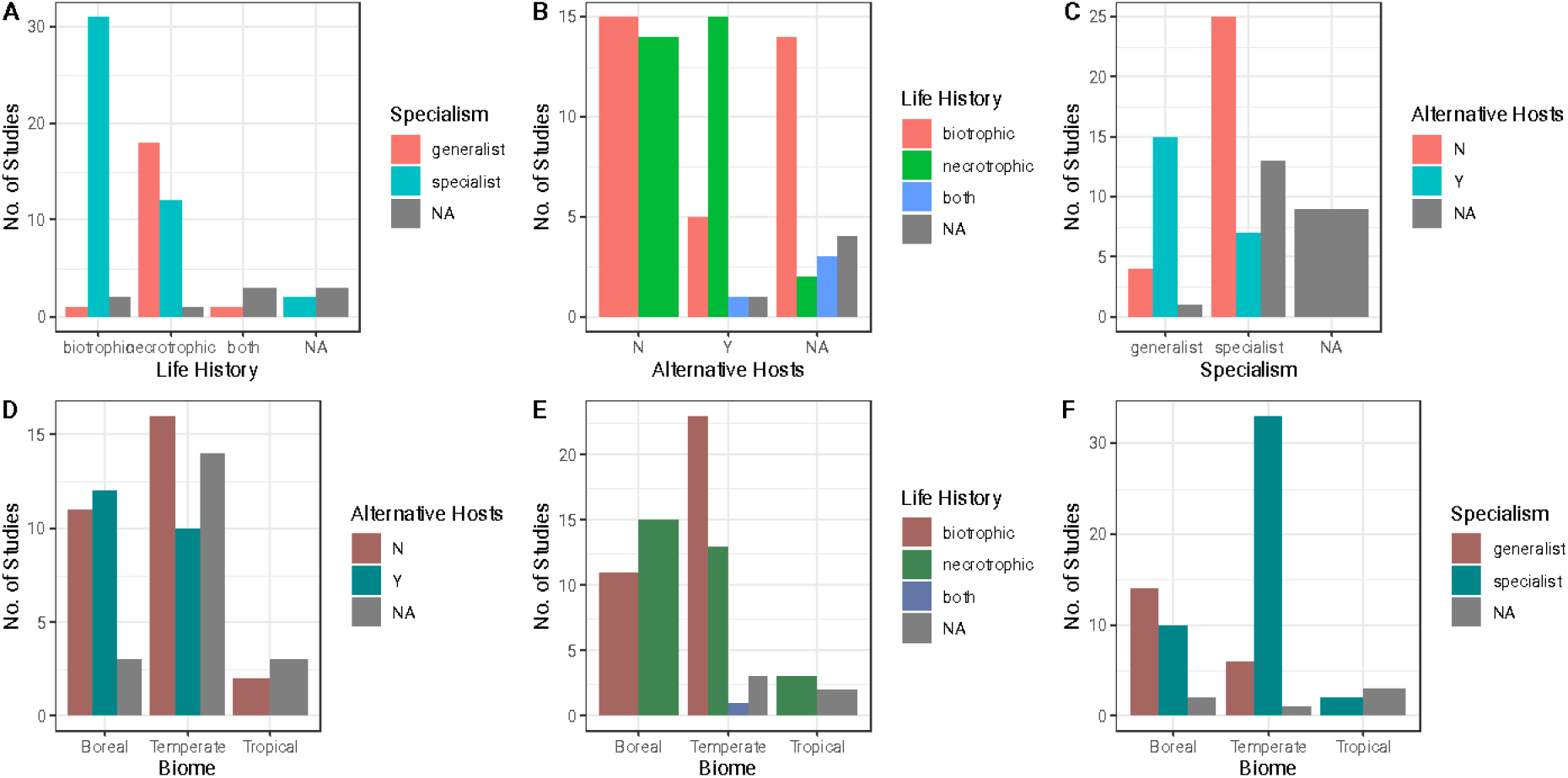
Bar charts showing categorical moderators used in mixed effects meta-analysis models, plotted against number of studies. Shown in each plot are the numbers of studies in each category, for two categorical moderators. Several moderators are well confounded (A, C, E, F). We therefore chose to analyse moderators separately rather than including them in the same models. Total no. of studies *(k)* = 74.

**Figure S2:**
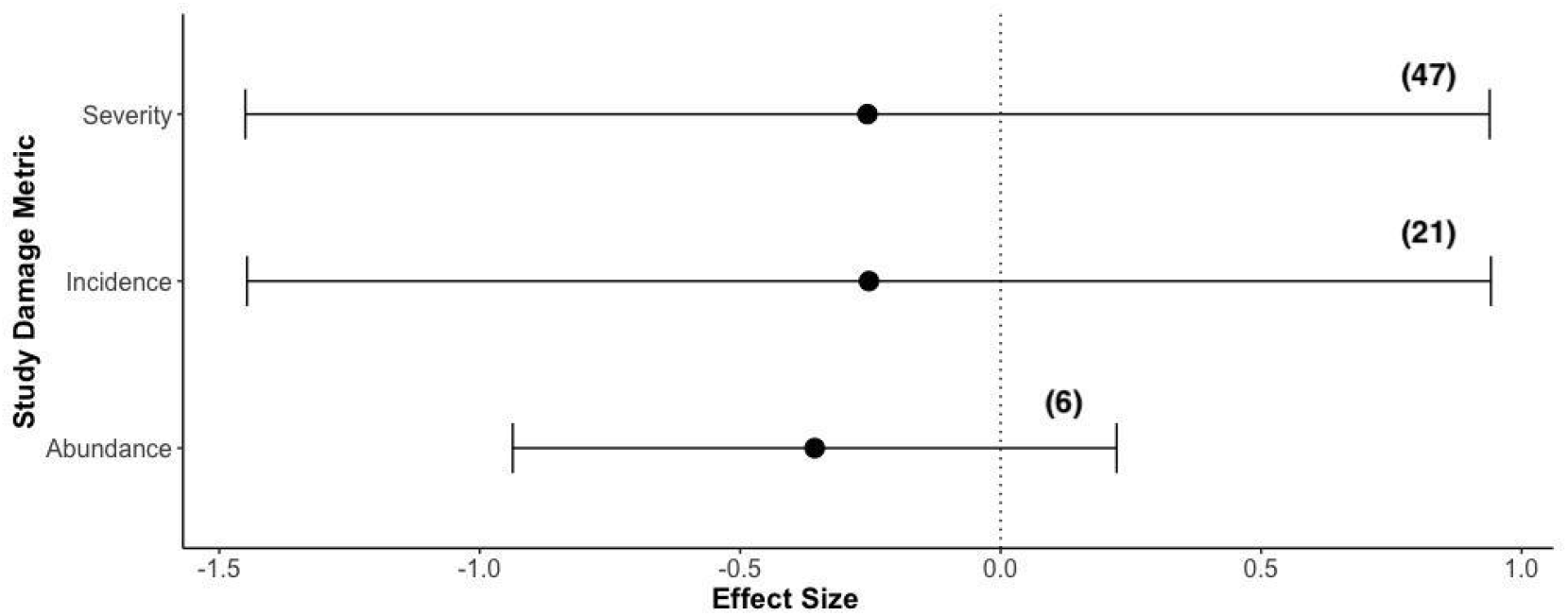
Effect of diversity metric used in studies on effect size (Hedges’ *d*). Shown is the mean effect size +/- 95% confidence intervals for each group. Metrics of pathogen damage fell into two main categories: pathogen incidence (*k =* 21 studies) and pathogen severity (*k =* 47 studies), with a smaller number (*k =* 6 studies) that assessed pathogen abundance. Diversity metric was a non-significant predictor of effect size Q_M_ = 0.11, *p* = 0.95, *df* = 2). In brackets is the number of studies in each group (total *k* = 72 studies).

**Figure S3:**
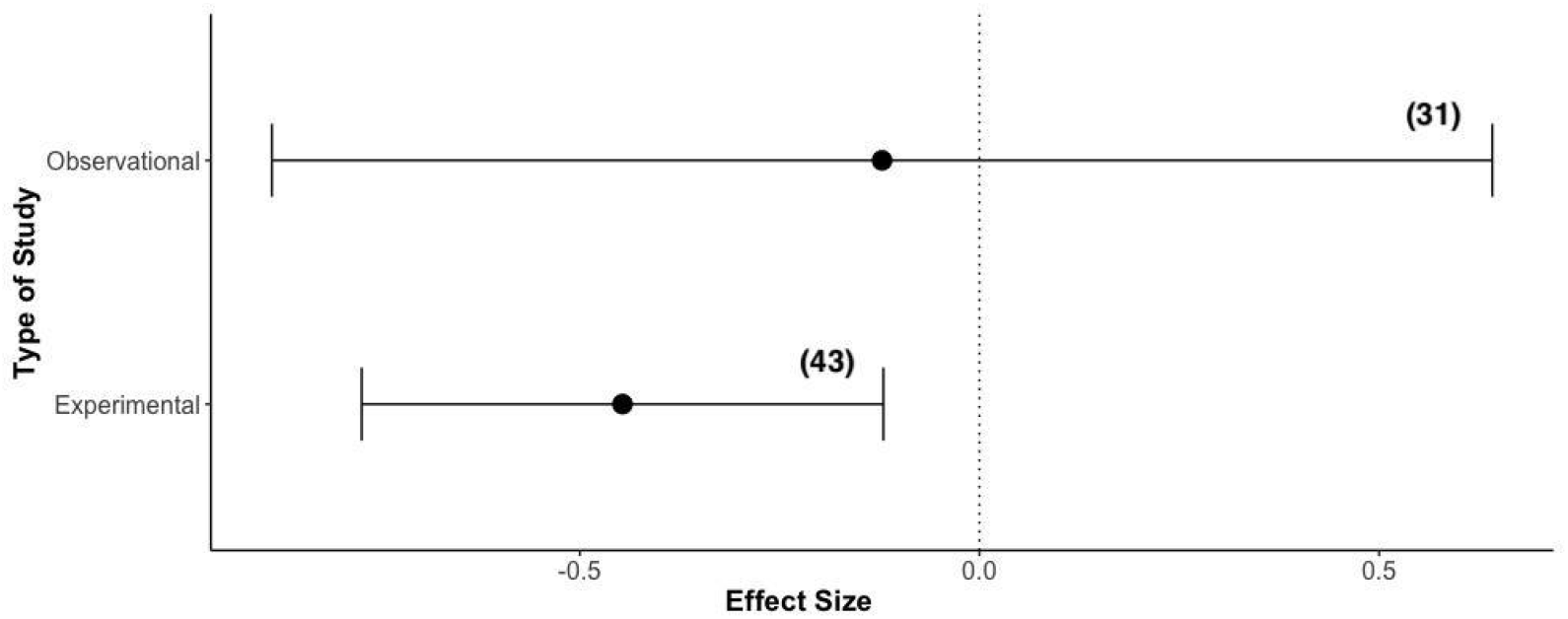
Effect of study type (experimental versus observational) on effect size (Hedges’ *d*). Shown is the mean effect size +/- 95% confidence intervals for each group. Study type was a non-significant predictor of effect size Q_M_ = 2.12, *p* = 0.14, *df* = 1). Damage in experimental studies was significantly lower in mixed forests compared to monocultures, but the effect was not significantly different from observational studies (Fig. 6). In brackets is the number of studies in each group (total *k* = 72 studies).

**Figure S4:**
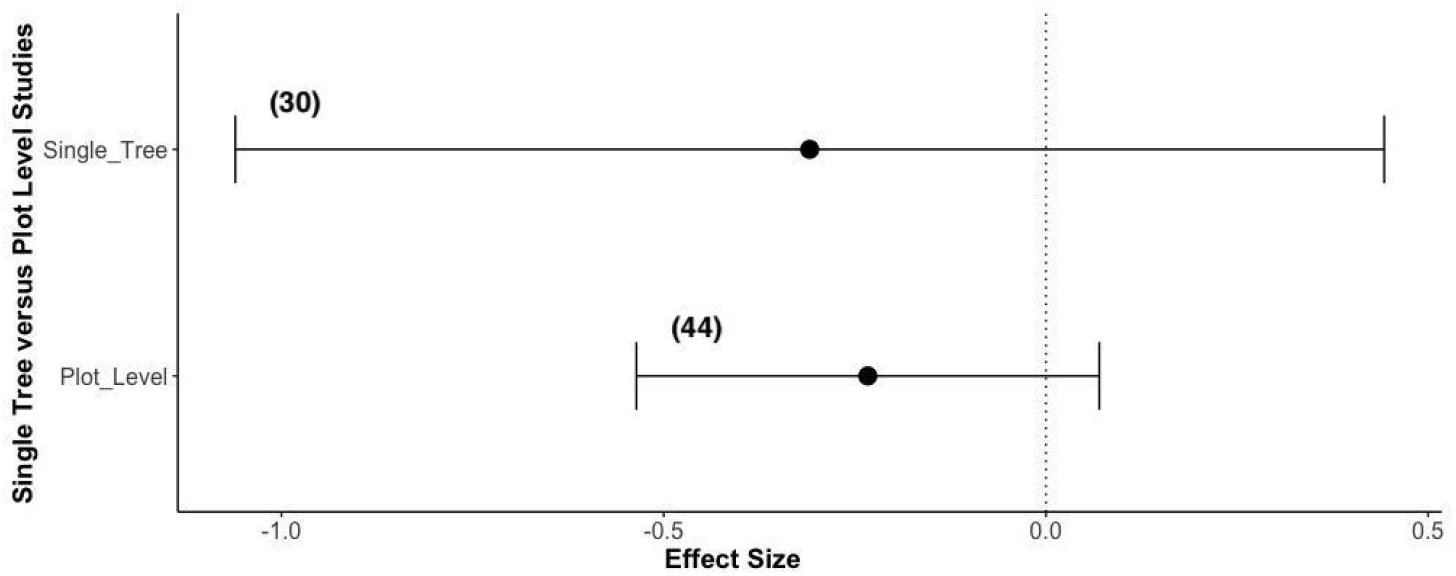
Effect of study level (single tree level versus plot level estimates) on effect size (Hedges’ *d*). Shown is the mean effect size +/- 95% confidence intervals for each group. Study type was a non-significant predictor of effect size Q_M_ = 0.11, *p* = 0.74, *df* = 1). In brackets is the number of studies in each group (total *k* = 72 studies).

**Figure S5:**
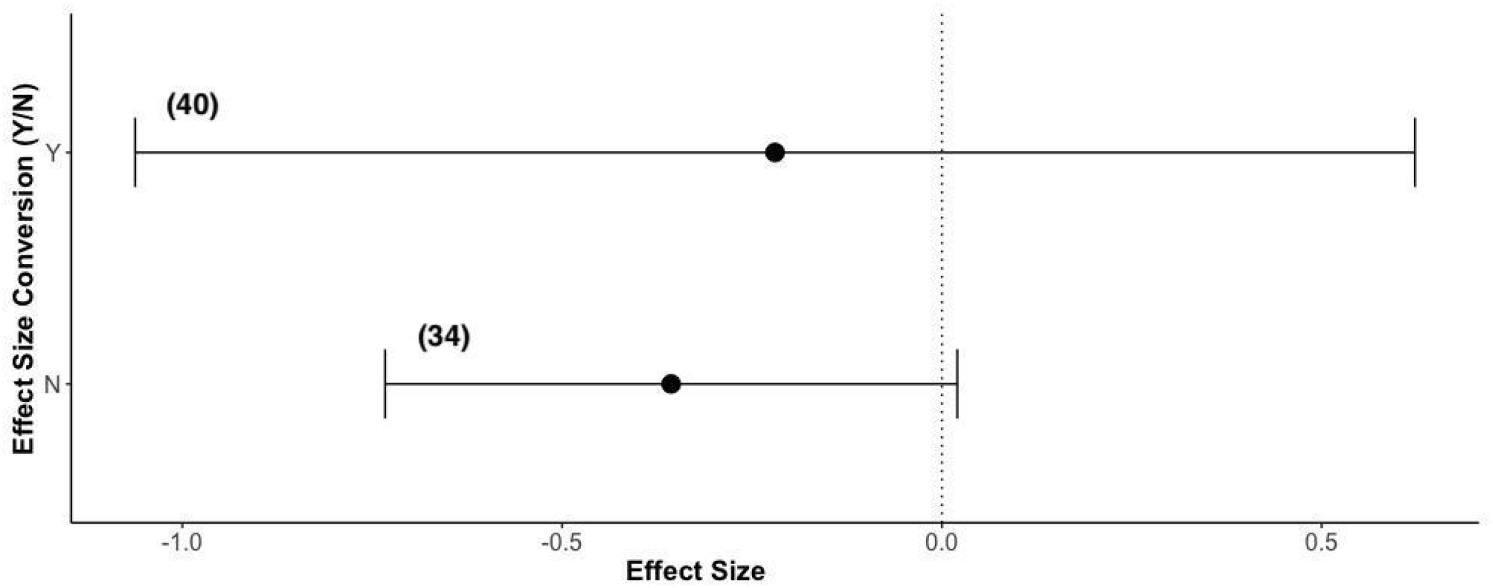
Comparison of directly calculated effect sizes and those converted from other metrics of effect size. Conversions were mostly from correlation coefficients, in some cases *p-*values and χ2 statistics. Shown is the mean effect size +/- 95% confidence intervals for each group. Effect size conversion was a non-significant predictor of effect size Q_M_ = 0.33, *p* = 0.57, *df* = 1). In brackets is the number of studies in each group (total *k* = 72 studies).

**Figure S6.**
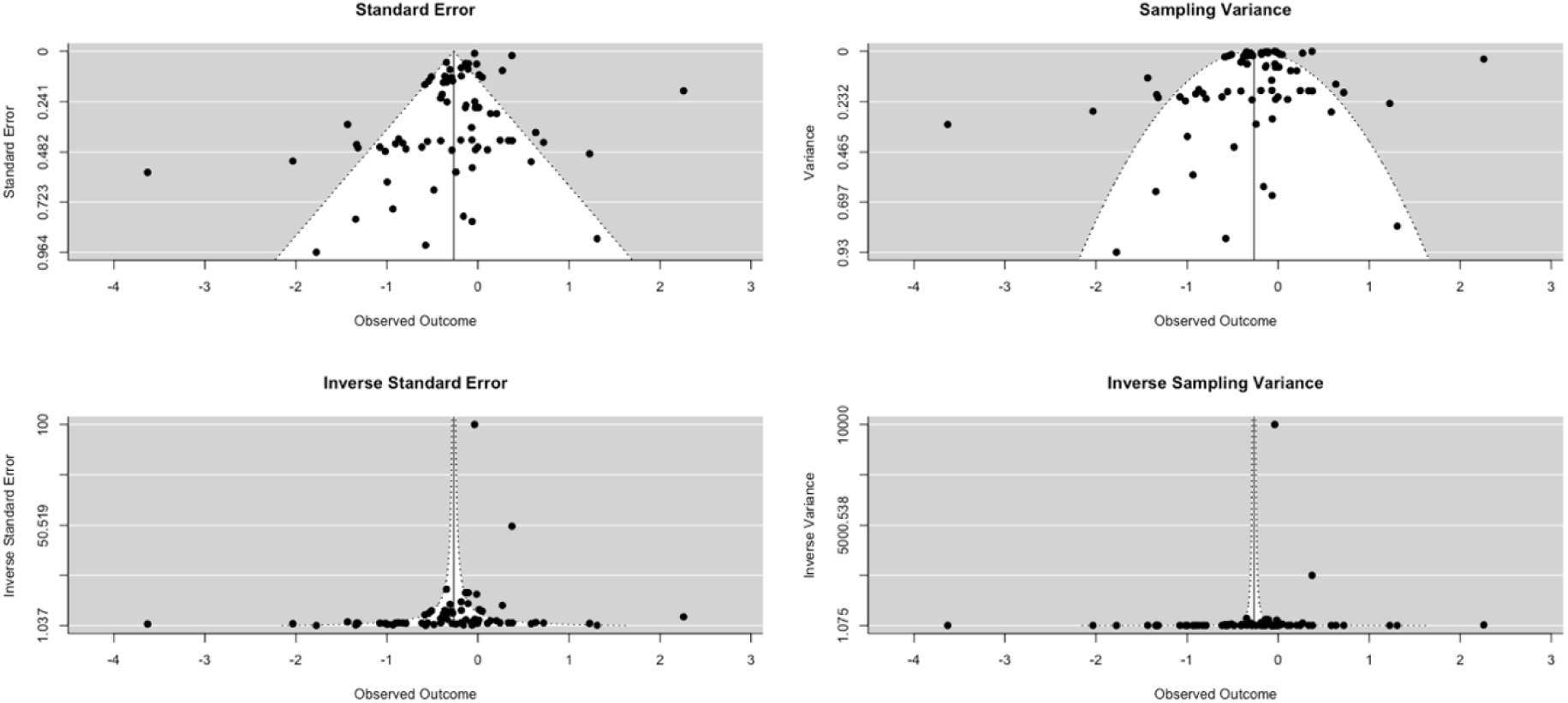
Funnel plots showing observed outcome (study effect sizes) against study standard error, variance, and inverse standard error/inverse variance. Plotted are the effect sizes. The black vertical line indicates the overall effect size, with the white region showing the boundaries within which studies would lie given perfect symmetry.

