## Supplementary Materials for "Tree diversity reduces pathogen damage in temperate forests: a systematic review and meta-analysis"

| PECO question component | Terms | Search terms |
| --- | --- | --- |
| Population | Forest | tree* OR forest* OR plantation OR wood* OR orchard OR coppice OR shrub OR abies OR acer OR betula OR eucalyptus OR fagus OR fraxinus OR picea OR pinus OR populus OR quercus |
| Exposure | Mixed Stands | "mixed species" OR "mixed-species" OR "mixed stand*" OR "tree divers*" OR "tree species divers*" OR "host species divers*" OR "host divers*" OR "community divers*" OR “forest divers*” OR "forest species divers*" OR "divers* stand" OR "divers* plant*" OR "divers* forest" OR "divers* wood*" OR "tree species rich*" OR "host species rich*" OR "forest species rich*" OR polyculture* OR "landscape heterogen*" OR "heterogen* landscape" OR "species mix*" OR "diversity-disease" OR "species composition" |
| Comparator | Forest Pathogen Damage | pathogen OR disease OR infection OR "root rot" OR "butt rot" OR fung* OR lesion OR canker OR bacteria* OR viral OR virus OR Phytophthora OR Armillaria OR Heterobasidion |

**Table S1:** List of search strings used in bibliographic databases and their relationship to each component of the primary question. Each section was combined using ‘AND’ (for exact search strings with Boolean operators used in each database, see Table S2).

| **Database** | **Search string used** | **Number of hits** | **Date** |
| --- | --- | --- | --- |
| Scopus | ( TITLE-ABS-KEY ( ( tree* OR forest* OR plantation OR wood* OR orchard OR coppice OR shrub OR abies OR acer OR betula OR eucalyptus OR fagus OR fraxinus OR picea OR pinus OR populus OR quercus ) AND ( "mixed species" OR "mixed-species" OR "mixed stand*" OR "tree divers*" OR “forest divers*” OR "tree species divers*" OR "host species divers*" OR "host divers*" OR "community divers*" OR "forest species divers*" OR "divers* stand" OR "divers* plant*" OR "divers* forest" OR "divers* wood*" OR "tree species rich*" OR "host species rich*" OR "forest species rich*" OR polyculture* OR "landscape heterogen*" OR "heterogen* landscape" OR "species mix*" OR "diversity-disease" OR "species composition" ) AND ( pathogen OR disease OR infection OR "root rot" OR "butt rot" OR fung* OR lesion OR canker OR bacteria* OR viral OR virus OR phytophthora OR armillaria OR heterobasidion ) ) ) AND ( LIMIT-TO ( SUBJAREA , "AGRI " ) OR LIMIT-TO ( SUBJAREA , " ENVI " ) ) | 1,744 (limited to Topics: Agricultural/Environmental Science and Biological Sciences) | 13^th^ December 2017 |
| Web of Science (All databases: WoS Core Collection 1945-present, BioSIS Citation Index, Current Contents Connect, Data Citation Index, Derwent Innovations Index, Inspec, KCI-Korean Journal Database, MEDLINE, Russian Science Citation Index, SciELO Citation Index, Zoological Record) | TS = (tree* or forest* or plantation or wood* or orchard or coppice or shrub or abies or acer or betula or eucalyptus or fagus or fraxinus or picea or pinus or populus or quercus) AND TS = ("mixed species" or "mixed-species" or "mixed stand*" or "tree divers*" or "tree species divers*" or “forest divers*” or "host species divers*" or "host divers*" or "community divers*" or "forest species divers*" or "divers* stand" or "divers* plant*" or "divers* forest" or "divers* wood*" or "tree species rich*" or "host species rich*" or "forest species rich*" or polyculture* or "landscape heterogen*" or "heterogen* landscape" or "species mix*" or "diversity-disease" or "species composition") AND TS = (pathogen or disease or infection or "root rot" or "butt rot" or fung* or lesion or canker or bacteria* or viral or virus or Phytophthora or Armillaria or Heterobasidion) | 5,703 (limited to Research Areas within Science and Technology: (Agriculture OR Forestry OR Environmental Sciences OR Ecology OR Plant sciences) | 13^th^ December 2017 |
| CAB Abstracts (Database: 1910 – 2017) | (tree* or forest* or plantation or wood* or orchard or coppice or shrub or abies or acer or betula or eucalyptus or fagus or fraxinus or picea or pinus or populus or quercus) AND ("mixed species" or "mixed-species" or "mixed stand*" or (mixed adj1 stand) or "tree divers*" or "tree species divers*" or "host species divers*" or "host divers*" or "community divers*" or "forest divers*" or "forest species divers*" or "tree species divers*" or (forest adj1 divers*) or (tree adj1 divers*) or "divers* stand" or "divers* plant*" or "divers* forest" or "divers* wood*" or "tree species rich*" or "host species rich*" or "forest species rich*" or polyculture* or "landscape heterogen*" or "heterogen* landscape" or "species mix*" or "diversity-disease" or "species composition") AND (pathogen or disease or infection or "root rot" or "butt rot" or fung* or lesion or canker or bacteria* or viral or virus or Phytophthora or Armillaria or Heterobasidion) | 2,756  Searching abstract, title, original title, broad terms, heading words, identifiers, cabicodes | 13^th^ December 2017 |

**Table S2:** List of search strings used in each major database queried during the systematic review, with number of hits returned and date initially searched.

NB: Google Scholar does not permit the use of Boolean operators, so search terms were modified and the first 1000 results downloaded from Google Scholar. We tried several search term combinations in Google Scholar to maximise the number of test library studies found while minimising duplication with the rest of the literature searches. Final search terms used in Google Scholar were: ‘ “mixed stand” forest pathogen ‘ (Search completed on 21^st^ February 2018).

### ***Appendix II: Test Library used in Scoping Searches***

### ***Appendix III: List of stakeholders consulted during the literature searching stage***

Centre for International Forest Research (CIFOR)

EU COST action on Mixed Forests (EUMIXFOR)

EUFORGEN

European Forest Institute

Food and Agriculture Organisation of the United Nations

Forest Europe

Forest Research (UK)

French National Institute for Agricultural Research (INRA)

International Union of Forestry Research Organisations (IUFRO)

United States Department of Agriculture

World Agroforestry Centre

### ***Appendix IV: Metadata Form sent to stakeholders from TreeDivNet contributing unpublished data sets***

| TreeDivNet Pathogen Data for Systematic Synthesis | |
| --- | --- |
| Name: | Institution: |
| Position: | Contact email: |
| Contact address: | Experiment name: |
| Location of experiment (latitude and longitude): | Dates of data collection: |
| Year planted: |  |
| Who collected the data? | Contact email of the person who collected the data (if different from above): |
| Tree species planted in experiment: | Tree species sampled for pathogens: |
| Please provide a brief description of sampling protocol, including the number of plots sampled and treatment levels, number of trees sampled per plot, number of leaves/shoots sampled per tree. | |
| How were pathogens identified (e.g. by eye in the field, microscopically from sampled leaves, molecular methods)? | |
| What is the metric of pathogen damage recorded? (e.g. % damage per leaf, number of stem lesions, etc). | |
| Further comments | |

### ***Appendix V: Supplementary Figures***

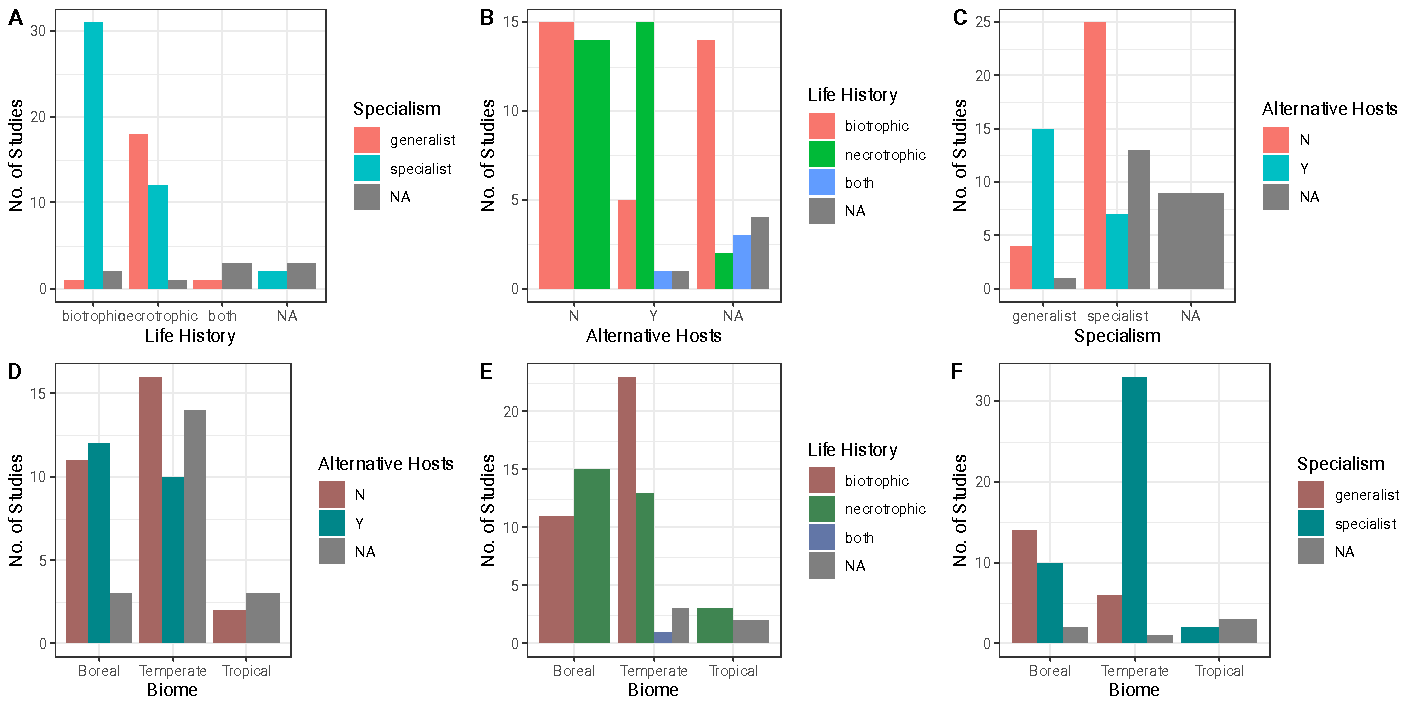

**Figure S1.** Bar charts showing categorical moderators used in mixed effects meta-analysis models, plotted against number of studies. Shown in each plot are the numbers of studies in each category, for two categorical moderators. Several moderators are well confounded (A, C, E, F). We therefore chose to analyse moderators separately rather than including them in the same models. Total no. of studies *(k)* = 74.

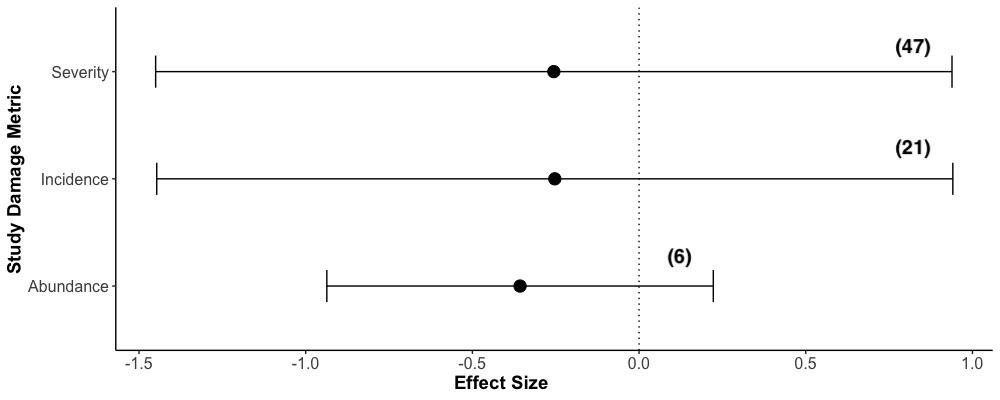

**Figure S2:** Effect of diversity metric used in studies on effect size (Hedges’ *d*). Shown is the mean effect size +/- 95% confidence intervals for each group. Metrics of pathogen damage fell into two main categories: pathogen incidence (*k =* 21 studies) and pathogen severity (*k =* 47 studies), with a smaller number (*k =* 6 studies) that assessed pathogen abundance. Diversity metric was a non-significant predictor of effect size Q_M_ = 0.11, *p* = 0.95, *df* = 2). In brackets is the number of studies in each group (total *k* = 72 studies).

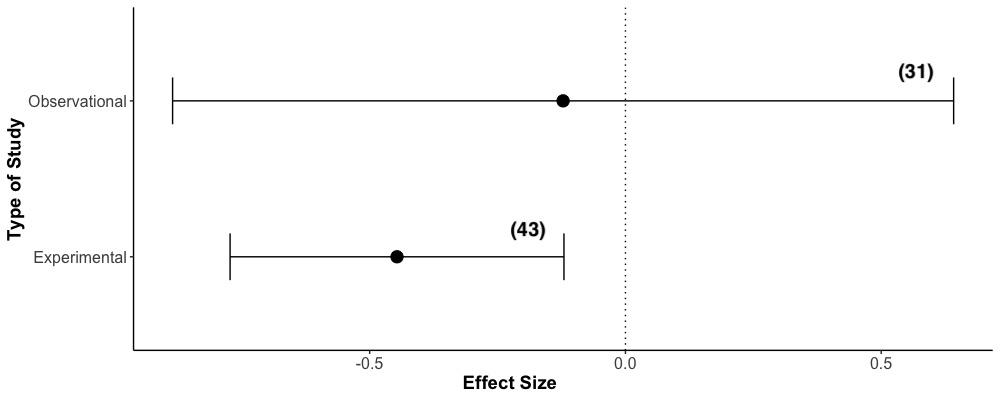

**Figure S3:** Effect of study type (experimental versus observational) on effect size (Hedges’ *d*). Shown is the mean effect size +/- 95% confidence intervals for each group. Study type was a non-significant predictor of effect size Q_M_ = 2.12, *p* = 0.14, *df* = 1). Damage in experimental studies was significantly lower in mixed forests compared to monocultures, but the effect was not significantly different from observational studies (Fig. 6). In brackets is the number of studies in each group (total *k* = 72 studies).

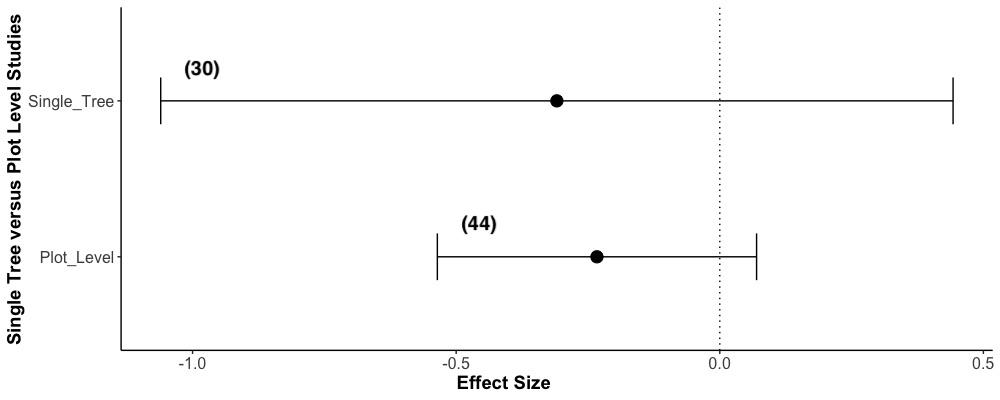

**Figure S4:** Effect of study level (single tree level versus plot level estimates) on effect size (Hedges’ *d*). Shown is the mean effect size +/- 95% confidence intervals for each group. Study type was a non-significant predictor of effect size Q_M_ = 0.11, *p* = 0.74, *df* = 1). In brackets is the number of studies in each group (total *k* = 72 studies).

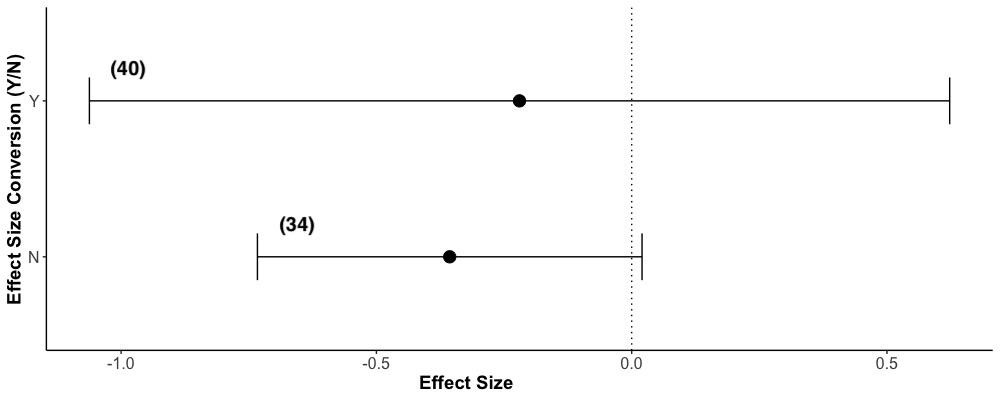

**Figure S5:** Comparison of directly calculated effect sizes and those converted from other metrics of effect size. Conversions were mostly from correlation coefficients, in some cases *p-*values and χ2 statistics. Shown is the mean effect size +/- 95% confidence intervals for each group. Effect size conversion was a non-significant predictor of effect size Q_M_ = 0.33, *p* = 0.57, *df* = 1). In brackets is the number of studies in each group (total *k* = 72 studies).

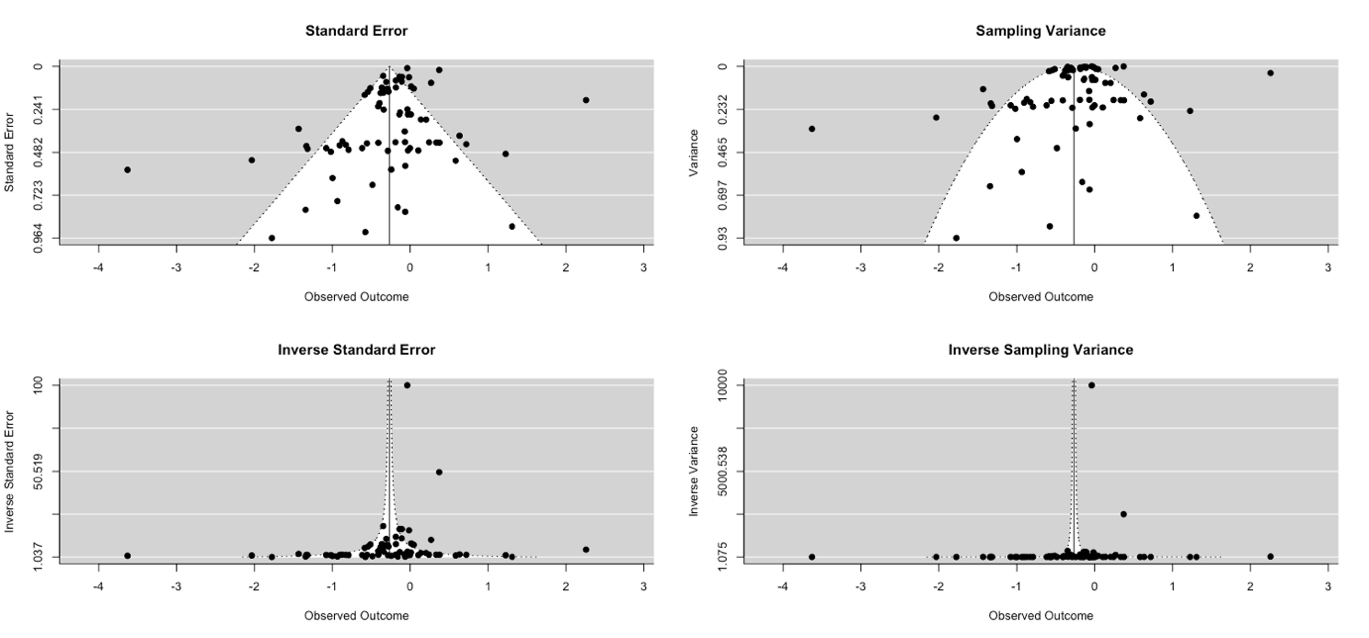

**Figure S6.** Funnel plots showing observed outcome (study effect sizes) against study standard error, variance, and inverse standard error/inverse variance. Plotted are the effect sizes. The black vertical line indicates the overall effect size, with the white region showing the boundaries within which studies would lie given perfect symmetry.
